# Oral dosing of the nucleoside analog obeldesivir is efficacious against RSV infection in African green monkeys

**DOI:** 10.1101/2025.02.24.639976

**Authors:** Jared Pitts, J. Lizbeth Reyes Zamora, Savrina Manhas, Thomas Aeschbacher, Josolyn Chan, Vincent Cutillas, Varsha Nair, Nicolas C. Riola, Arya Vijjapurapu, Meghan S. Vermillion, Stacey Eng, Christopher Richards, Dong Han, Jason K. Perry, Subhra Chaudhuri, Iris Liu, Clarissa Martinez, Nadine Peinovich, Kai-Hui Sun, Arthur Cai, Ross Martin, Jasmine Moshiri, Charlotte Hedskog, Darius Babusis, Dustin S. Siegel, Rao Kalla, Vasanthi Avadhanula, Pedro A. Piedra, Kim Stobbelaar, Peter L. Delputte, Caleb Marceau, Roberto Mateo, Jenny Svarovskaia, Hongmei Mo, Raju Subramanian, Richard L. Mackman, Tomas Cihlar, Simon P. Fletcher, John P. Bilello

**Author notes:** These authors contributed equally to this work (Jared Pitts, J. Lizbeth Reyes Zamora). Corresponding author Gilead Sciences, Inc., 333 Lakeside Drive, Foster City, CA 94404.

## Abstract

Respiratory syncytial virus (RSV) is a significant cause of morbidity and mortality in high-risk populations. Although prophylactic options are available, there are no effective oral therapeutics for RSV infection. Obeldesivir (ODV) is an orally bioavailable prodrug of the nucleoside analog GS-441524, which is converted intracellularly to its active nucleoside triphosphate and inhibits the RSV RNA polymerase. Here we report the potent antiviral activity of ODV against geographically and temporally diverse RSV A and B clinical isolates (EC_50_: 0.20-0.66 μM). Resistance selection studies with ODV and GS-441524 against RSV identified a single amino acid substitution, I777L, in the L polymerase with reduced susceptibility (3.3-3.8-fold) to ODV and GS-441524, indicating a high barrier for resistance development. In an African green monkey RSV infection model, once-daily oral ODV doses of 30 or 90 mg/kg initiated ∼24 hours post-infection significantly reduced log_10_ viral RNA copies/mL×day area under the curve by 69-92% in the upper and lower respiratory tracts. Together, these preclinical data support the clinical evaluation of ODV for the treatment of RSV infection.

## INTRODUCTION

Respiratory syncytial virus (RSV) is a negative-strand RNA virus that infects most of the population in early childhood^1^. In healthy individuals, RSV infection is primarily limited to the upper respiratory tract resulting in mild cold-like symptoms. However, in young children, older adults or adults with underlying comorbidities, RSV can cause lower respiratory tract infections leading to more severe outcomes such as bronchiolitis, pneumonia, and/or decreased lung function. Globally, there are an estimated 33 million pediatric RSV cases per year, resulting in ∼3 million hospitalizations and ∼100,000 deaths^2,3^. Individuals with cardiopulmonary comorbidities or who are immunocompromised are at higher risk of hospitalization and death resulting from RSV infection^4^. Despite the availability of effective prophylactic options for high-risk populations, there remains an unmet need for an oral treatment for RSV.

The RSV genome encodes 11 viral proteins, several of which have been explored as prophylactic and therapeutic targets. Fusion protein (F) inhibitors (e.g. presatovir, ziresovir), the nucleoprotein (N) inhibitor zelicapavir, and the large polymerase (L) inhibitor lumicitabine, have all demonstrated efficacy in human RSV challenge studies^5–9^. However, only ziresovir has subsequently shown signs of efficacy in clinical trials of naturally infected individuals^10^. Remdesivir (RDV, VEKLURY®), an intravenous (IV) administered phosphoramidate prodrug of the nucleoside GS-441524, exhibits antiviral activity against RSV, SARS-CoV-2, and other RNA viruses^11,12^. RDV is efficiently metabolized intracellularly to its active nucleoside triphosphate (NTP, GS-443902) which inhibits viral RNA-dependent RNA polymerases to limit viral replication^11^. Although RDV was the first antiviral approved for the treatment of hospitalized patients with COVID-19^13^, it was initially invented as a potent inhibitor of RSV, with prophylactic efficacy in an African green monkey (AGM) infection model of RSV^14^. However, the IV route of administration for RDV limits its utility for treatment of RSV in nonhospitalized individuals.

Orally administered RDV and the parent nucleoside, GS-441524, have limited oral bioavailability in non-human primates (<1% and <10%, respectively)^14^. RDV and GS-441524 both metabolize to the same active triphosphate, GS-443902, and thus share the same antiviral spectrum and mechanism of action. Obeldesivir (ODV; GS-5245) (Figure 1A) is a 5’-mono-isobutyryl ester prodrug of GS-441524 designed to increase the oral bioavailability of GS-441524^15^. Following oral administration, ODV is extensively cleaved pre-systemically to the parent nucleoside, achieving efficacious GS-441524 systemic exposures at lower doses than would be required with direct oral dosing of GS-441524. ODV in vivo antiviral efficacy is driven by high systemic levels of GS-441524, which has been demonstrated in non-human primate models of SARS-CoV-2, Sudan ebolavirus, and Marburg virus^15–17^.

**Figure 1.**
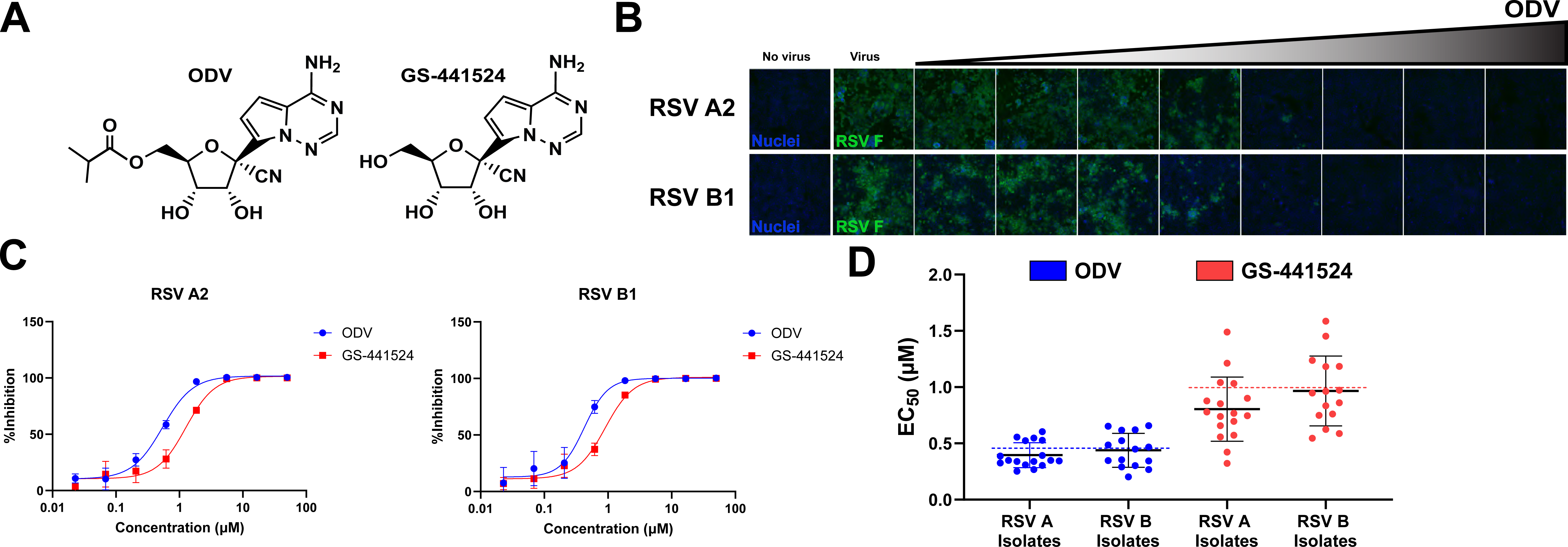
In vitro antiviral activity of ODV and its parent nucleoside against RSV laboratory strains and clinical isolates. (A) Structure of ODV and its systemic metabolite GS-441524 (parent nucleoside). (B) Representative images for ODV dose response against laboratory strains RSV A2 and RSV B1 in HEp-2 cells. RSV F protein detection in green and nuclei detection in blue. (C) Representative dose-response curves for ODV and GS-441524 against RSV A2 and B1. (D) Distribution of ODV (blue) and GS-441524 (red) half-maximal effective concentrations (EC_50_s) of RSV A clinical isolates (n=17) and RSV B clinical isolates (n=15). Black horizontal lines represent the mean RSV clinical isolate EC_50_ ± standard deviation. The mean RSV A2 EC_50_ for ODV (red) and GS-441524 (blue) are represented by the dashed lines.

In this study, we show that ODV has antiviral activity against diverse RSV clinical isolates and has a high barrier to resistance emergence in vitro. Further, oral treatment with ODV is efficacious in an AGM model of RSV infection. Supported by this preclinical data, oral ODV is currently being evaluated for the treatment of RSV infection in children and high-risk adults (NCT06784973, NCT06585150).

## RESULTS

### Obeldesivir and its parent nucleoside have potent antiviral activity against diverse RSV A and B isolates

Two genetically distinct subgroups of RSV (A and B) can cause severe disease and co-circulate in alternating annual seasonal patterns. Antiviral activity of ODV and the parent nucleoside GS-441524 was evaluated in HEp-2 cells against RSV A2 and B1 laboratory strains (LS) utilizing a quantitative cellomics-based antiviral assay which measures F expression through antibody binding to a conserved F antigenic site (Figure 1B and C)^18–20^. ODV and GS-441524 inhibited replication of RSV laboratory strains with half-maximal effective concentrations (EC_50_s) of 0.43-0.46 µM and 0.91-1.0 µM, respectively (Figure 1C; Supplementary Tables 1 and 2; Supplementary Figure 1). No cytotoxicity was observed with ODV or GS-441524 in HEp-2 cells at concentrations up to 50 µM and 100 µM, respectively (Supplementary Table 1)^14^.

Thirty-two RSV clinical isolates collected between 1987-2023 were obtained from the US, UK, and Belgium. Sequencing and phylogenic analysis determined the A and B clinical isolates were genetically distinct, spanning 18 unique lineages, with several isolates assigned to lineages currently in circulation (A.D.5.2, A.D.3, A.D.1, B.D.4.1.1, and B.D.E.1) (Supplementary Tables 1 and 2 and Supplementary Figure 2) (https://nextstrain.org/rsv). The mean antiviral activities of ODV and GS-441524 against these diverse clinical isolates ranged from 0.20-0.66 µM and 0.32–1.59 μM, respectively (Supplementary Tables 1 and 2, Supplementary Figure 1). ODV and GS-441524 EC_50_ values against all RSV A and B isolates tested were comparable to the RSV A2 laboratory strain (0.32 to 1.59-fold) (Figure 1D, Supplementary Tables 1 and 2, Supplementary Figure 1).

### ODV and GS-441524 have a high in vitro barrier to resistance in the RSV A2 laboratory strain

To identify genetic changes associated with reduced susceptibility to ODV or its parent nucleoside, RSV A2 was serially passaged in HEp-2 cells with DMSO (as a negative control), ODV, GS-441524, or the F-inhibitor presatovir, which was previously shown to rapidly select for highly resistant variants^19,21^. RSV A2 cytopathic effects and syncytia formation were observed at presatovir concentrations >100× EC_50_ within 18 days (Passage [P] 3), selecting for substitutions associated with presatovir resistance ^22,23^ (Figure 2A, Supplementary Tables 3 and 4). In contrast, dose-escalations of ODV and GS-441524 gradually progressed to a maximum of 32× and 10× EC_50_ respectively over 91 days (P13) (Figure 2A, Supplementary Table 3). At day 49 (P7) of ODV selection, an I777L substitution near the active site of the L polymerase emerged and was fixed in the population by the next passage (Figure 2A, Supplementary Table 3, Supplementary Figure 3A). In the GS-441524 selection, a mixed population containing L1453L/W within the capping domain of the L polymerase emerged after 36 days (P5) and remained as a mixture through day 91 (Figure 2A, Supplementary Table 3, Supplementary Figure 3A).

**Figure 2.**
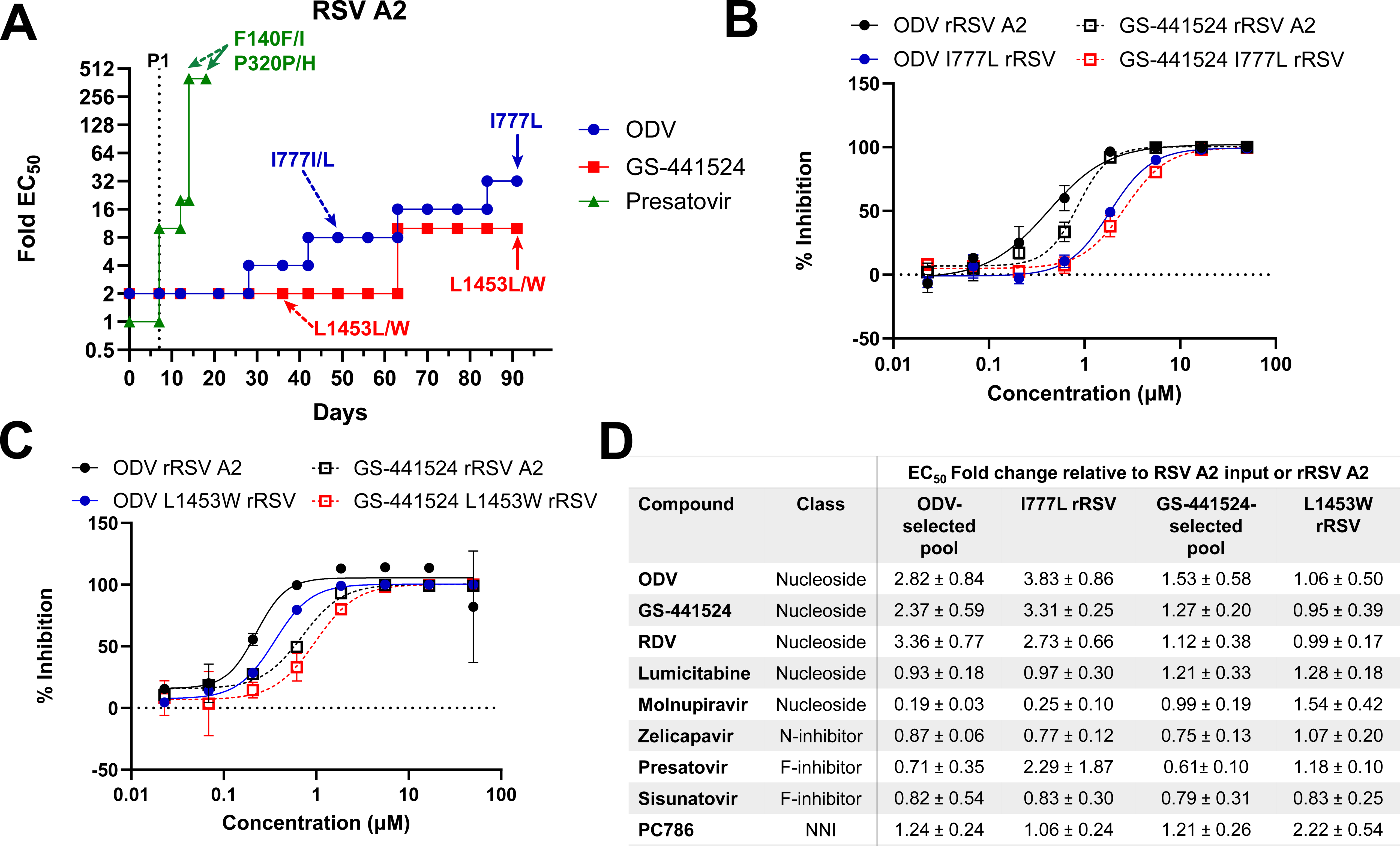
In vitro resistance selection of the RSV A2 laboratory strain and characterization of selected substitutions. (A) Progression of continuous passage of RSV A2 for 91 days under escalating doses of ODV, GS-441524, or presatovir. Substitutions in the L polymerase (ODV and GS-441524 selections) or the F protein (presatovir selection) were determined by Next Generation Sequencing (NGS) analysis. Substitutions at >15% prevalence and in ≥ 2 successive passages which were not identified in the DMSO controls are indicated by dashed arrows (first detection) and solid arrows (detection at final passage). (B) Representative dose-response curves of ODV and GS-441524 against I777L recombinant RSV (rRSV). (C) Representative dose-response curves of ODV and GS-441524 against L1453W rRSV. Wildtype rRSV A2 (black) served as a reference control for rRSV with an L substitution. (D) ODV- or GS-441524-selected pools and cloned rRSV A2 bearing the I777L or L1453W substitutions in the L protein were tested against a panel of RSV inhibitors. Fold changes in EC_50_ values from the wildtype RSV A2 control are shown (n=2-4); NNI (non-nucleoside inhibitor).

The ODV-selected pool (propagated from P13) as well as a cloned recombinant RSV A2 (rRSV) bearing the I777L L polymerase substitution had marginal reductions in susceptibility to ODV, GS-441524 and RDV (2.37-3.83-fold), compared to the respective wild-type RSV A2 controls (Figure 2B and D, Supplementary Tables 5 and 6). In contrast, the GS-441524 selected-pool (containing a mixed L1453L/W population) and rRSV expressing the L1453W substitution remained fully susceptible to both ODV and GS-441524 (Figure 2C and D, Supplementary Tables 5 and 6). Both the ODV- and GS-441524-selected pools as well as the I777L and L1453W recombinant viruses remained fully susceptible to inhibitors with other mechanisms of action (Figure 2D, Supplementary Tables 5 and 6).

### ODV and GS-441524 maintain a high barrier to resistance in RSV clinical isolates

RSV A2 is a laboratory strain originally collected in 1961 that has undergone extensive passaging in HEp-2 cells. To identify potential ODV and GS-441524 resistance-associated substitutions in more recent RSV isolates, resistance selections were performed with RSV clinical isolates from hospitalized US pediatric patients from the 2004-2005 RSV season. RSV A clinical isolate HRSV/A/Texas.USA/79254/2004 and RSV B clinical isolate HRSV/B/Texas.USA/79362/2005 were serially passaged with ODV, GS-441524, or presatovir as described for RSV A2. The RSV A and B clinical isolates replicated efficiently at >100× EC_50_ presatovir concentrations by days 12 (P2) and 24 (P4), respectively, rapidly selecting known resistance-associated substitutions (Figure 3A and B, Supplementary Table 3)^22,23^. Presatovir-selected pools from the RSV A and B clinical selections were highly resistant to fusion inhibitors but not inhibitors targeting the N protein or the L polymerase (Supplementary Table 4). In contrast, the RSV A clinical isolate selection with ODV and GS-441524 did not progress beyond 8× or 4× EC_50_, respectively (Figure 3A). A mixed K1188K/E substitution in the L polymerase, distant from its active site, was transiently observed in the ODV selection; however, no substitutions were detected in the final passages (P13) of selection with ODV or GS-441524 (Supplementary Table 3, Supplementary Figure 3B). RSV B clinical isolate selections with ODV and GS-441524 did not exceed 10× and 5× EC_50_, respectively (Figure 3B). While mixed populations of S169S/P, T660T/A, C319C/R and C1881C/Y were observed in the L polymerase after 4-5 weeks (P4-5) of selection with GS-441524, no substitutions were detected during the 91 days of ODV selection (Figure 3B, Supplementary Table 3). Only the C319R substitution persisted in the GS-441524-selected RSV B virus pool through the end of selection (day 91). Structural analysis revealed that S169P, C319R, and C1881Y are distant from the active site and unlikely to confer resistance to ODV or GS-441524, whereas T660A is near the active site but was only transiently observed (Supplementary Table 3, Supplementary Figure 3C). Although prolonged passage of RSV B in GS-441524 selected for the C319R substitution, virus containing C319R from the last passages could not be rescued. We therefore evaluated rRSV expressing C319R against the same panel of inhibitors as for RSV A2. Consistent with the distant location of C319R from the L polymerase active site, C319R rRSV remained fully susceptible to ODV and GS-441524 (Figure 3C and D, Supplementary Table 6). Additionally, inhibitors with other mechanisms of action remained fully active against rRSV containing the C319R substitution (Figure 3D, Supplementary Table 6).

**Figure 3.**
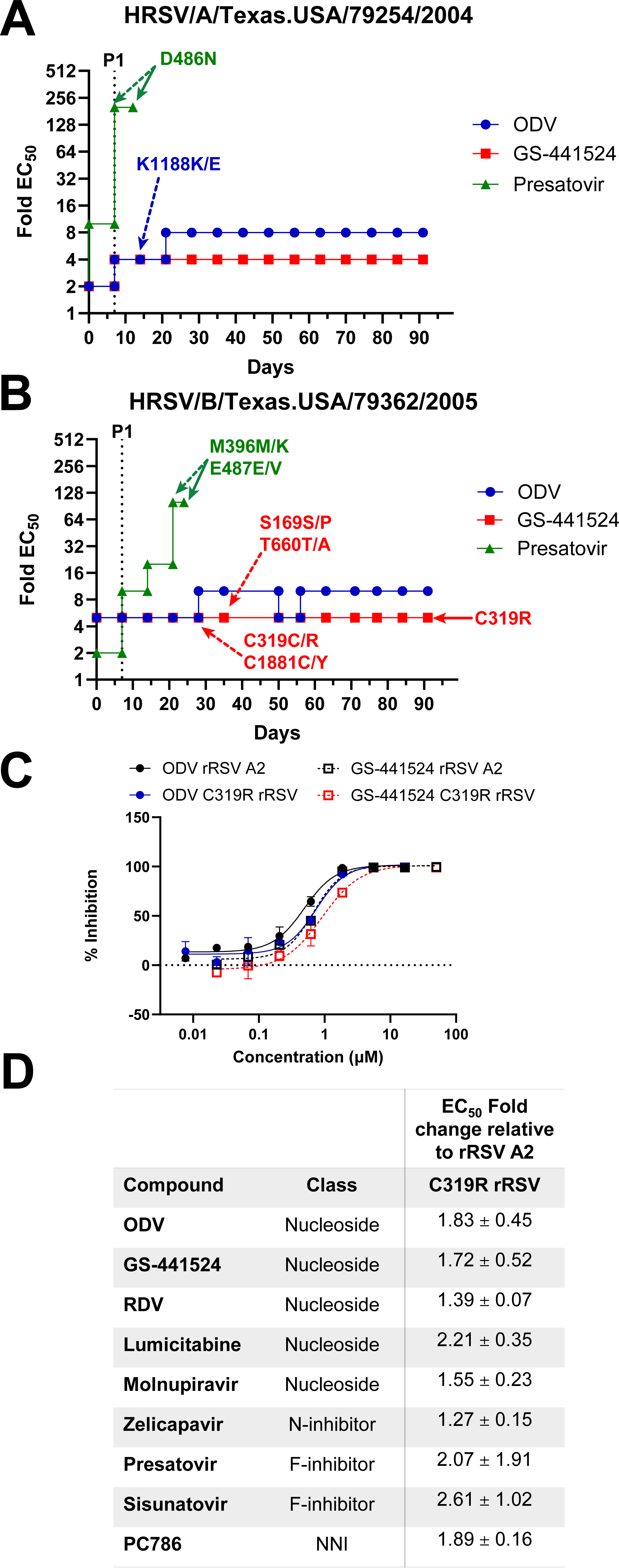
In vitro resistance selection of RSV A and B clinical isolates and characterization of the selected C319R substitution. Progression of continuous passage of (A) RSV A clinical isolate HRSV/A/Texas.USA/79254/2004 or (B) RSV B clinical isolate HRSV/B/Texas.USA/79362/2005 for 91 days under escalating doses of ODV, GS-441524, or presatovir. Substitutions in the L polymerase (ODV and GS-441524 selections) or the F protein (presatovir selection) were determined following NGS analysis. Substitutions at >15% prevalence in ≥ 2 successive passages and not identified in DMSO controls are indicated by dashed arrows (first detection) and solid arrows (detection at final passage). (C) Representative dose-response curves of ODV and GS-441524 against recombinant RSV (rRSV) A2 bearing C319R in the L polymerase. Reference wildtype rRSV A2 (black) was tested concurrently with C319R rRSV. (D) Cloned rRSV A2 bearing the C319R was tested against a panel of RSV inhibitors. Fold changes in EC_50_ values from the wildtype rRSV A2 control are shown (n=2); NNI (non-nucleoside inhibitor).

### Therapeutic efficacy of ODV against RSV in African green monkeys

African green monkeys (*Chlorocebus aethiops*) are semi-permissive to RSV infection, having comparable viral replication in both the upper and lower respiratory tracts to experimentally infected humans, but accompanied by limited symptomology^24,25^. This model has previously been utilized to assess the efficacy of several RSV vaccines and antiviral therapeutics^14,26–28^. To evaluate the efficacy of ODV against RSV, a cohort of 15 AGMs were infected with RSV A2 via intranasal and intratracheal instillation. Daily oral administration of ODV at 30 or 90 mg/kg was initiated 20-28 hours post-infection and continued for a total of 6 doses. Viral RNA levels in the upper and lower respiratory tracts were measured from throat swabs and bronchoalveolar lavage fluid (BALF) collected every other day through 15 days post-infection (Figure 4A).

**Figure 4.**
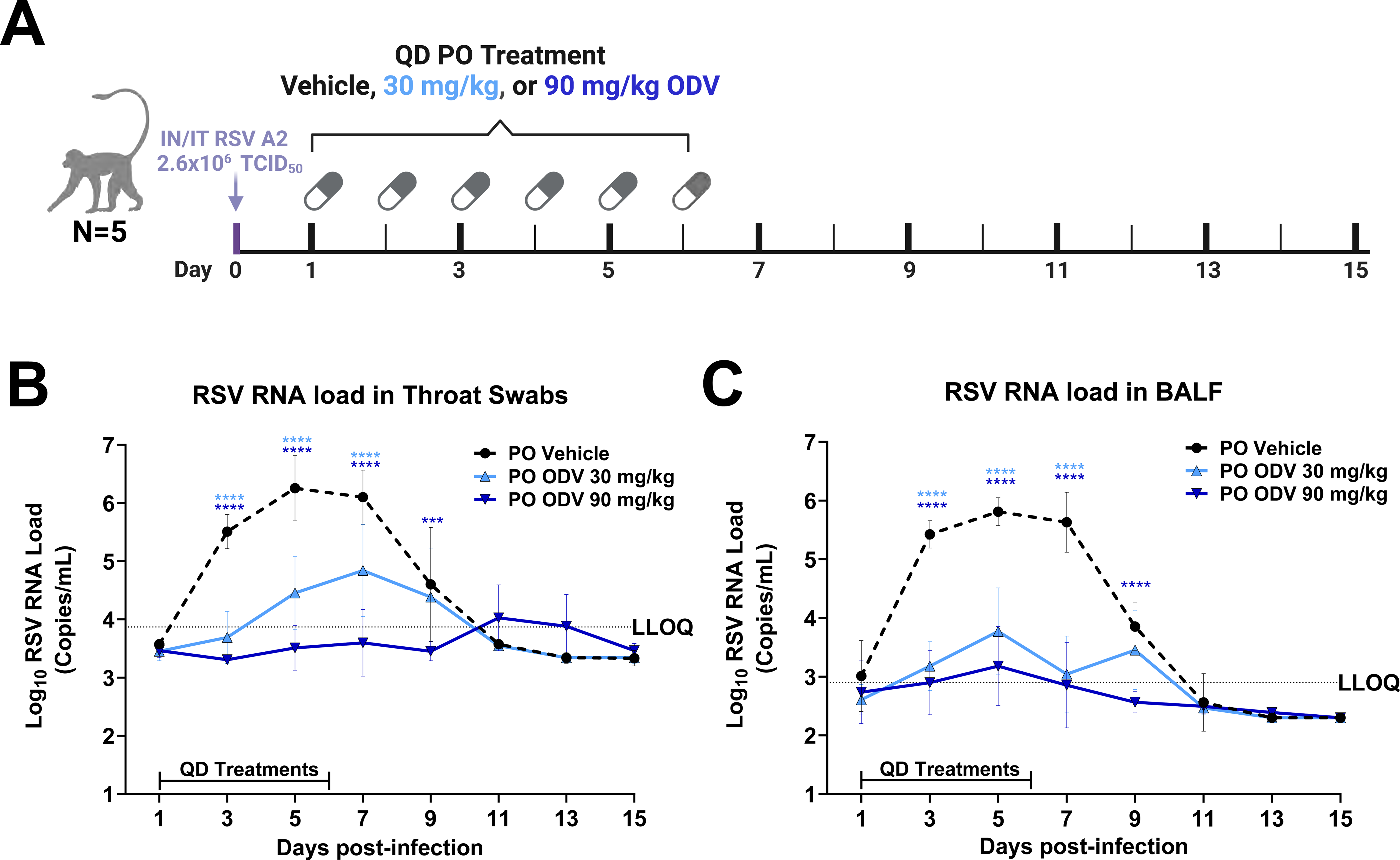
Efficacy of oral ODV in RSV infected African green monkeys. (A) Schematic of study design for AGM study. Animals (n=5 per group) were infected with ∼2.6 x 10^6^ TCID_50_ RSV A2 by intranasal/intratracheal instillation. Starting 20-28 hours post-infection, the animals were dosed with vehicle, 30 mg/kg ODV, or 90 mg/kg ODV once daily through day 6 post-infection (total of 6 doses). Bolded days represent timepoints at which throat swabs and BALF were collected for analysis. RSV RNA from throat swabs (B) or BALF samples (C) as determined by RT-qPCR for vehicle control (black dashed line), 30 mg/kg ODV (light blue line), and 90 mg/kg ODV (dark blue line). Samples with undetectable RNA levels were assigned a value of one quarter the lower limit of quantification (LLOQ), and samples with detectable RNA levels below the LLOQ were assigned values of one half the LLOQ before log_10_ transformation for analysis. The LLOQs are 3.3 and 2.3 log_10_ RNA copies/mL for the throat swab and BALF, respectively. The plots display the mean and standard deviation of the log_10_ transformed values. Statistical significance relative to the vehicle control was calculated using a two-way ANOVA with Bonferroni post hoc correction for multiple comparisons. Only statistically significant (p<0.05) differences are shown. \*\*\**p*<0.001. **** *p*<0.0001.

Compared to animals receiving vehicle, significant viral RNA reductions were observed in the upper respiratory tract following 30 mg/kg ODV treatment on days 3-7 post-infection, while ODV at 90 mg/kg significantly reduced viral RNA from days 3-9 post-infection (Figure 4B, Supplementary Table 7). At peak viral load (day 5), mean RSV RNA levels were reduced in a dose-dependent manner by 1.80 and 2.75 log_10_ copies/mL compared to the vehicle control in the 30 and 90 mg/kg groups, respectively. Consistent with these changes in viral load in the upper respiratory tract on individual days, 30 and 90 mg/kg ODV treatment significantly reduced the mean 15-day log_10_ RSV RNA area under the curve (AUC) relative to the vehicle control group (69% and 92%, respectively) (Supplementary Table 7).

ODV treatment also reduced viral RNA levels in the lower respiratory tract. The percent reduction in mean log_10_ viral RNA AUC in BALF samples for the ODV 30 and 90 mg/kg groups compared to the vehicle group were 78% and 90%, respectively. ODV treatment in both dose groups resulted in >2.0 log_10_ mean viral RNA levels relative to vehicle control at peak viral RNA load (day 5). Furthermore, 30 mg/kg and 90 mg/kg ODV significantly reduced viral RNA load from days 3-7 and days 3-9 post-infection, respectively, compared to vehicle (Figure 4C, Supplementary Table 7). Taken together, these results demonstrate that daily oral ODV treatment administered 20-28 hours post-infection is efficacious in a nonhuman primate model of RSV.

## DISCUSSION

RSV remains a major cause of respiratory disease for several high-risk populations with no oral therapeutic options available^29,30^. Several small molecule RSV antivirals such as zelicapavir, ziresovir, EDP-323 and S-337395 are currently in various stages of clinical evaluation. While multiple molecules have shown efficacy in human challenge studies ^5,21,31^, only the F-inhibitor ziresovir has shown signs of efficacy against natural RSV infection in a high-risk population^10^. Further evaluation of these clinical-stage drugs, as well as novel therapies with separate mechanisms of action, are needed to address the unmet disease burden of RSV.

Oral ODV was evaluated in Phase 3 COVID-19 clinical trials and was safe and well-tolerated, and significantly reduced viral RNA and infectious virus in nasal swabs compared to the placebo control^32–35^. The GS-441524 nucleoside metabolite in circulation following oral ODV administration was previously found to be potent against RSV, and the resulting NTP that is formed intracellularly is a potent inhibitor of the RSV polymerase^11,14,36^. In this report, we presented the preclinical profile of ODV which supported its clinical evaluation as an RSV therapeutic.

ODV and its parent nucleoside, GS-441524, had potent antiviral activity against RSV A and B laboratory strains as well as a comprehensive and divergent set of RSV A and B clinical isolates, including those observed in circulation as recently as 2023. In vitro resistance studies revealed that ODV and its parent nucleoside have a high barrier to resistance development, with substantially slower emergence of mutations in the L polymerase gene and only marginal changes in the susceptibility of selected variants compared to RSV isolates selected in the presence of the fusion inhibitor presatovir. Apart from the I777L substitution, long-term treatment of the RSV A2 laboratory strain and relevant RSV clinical isolates with ODV or its parent nucleoside selected substitutions in the L polymerase that were transient or remained fully susceptible to both compounds. The I777L substitution only emerged in the ODV selection with the RSV A2 laboratory strain, requiring more than 7 weeks to appear, and conferred <4-fold loss in susceptibility to ODV and GS-441524. Bioinformatic analysis of publicly available RSV sequences identified no polymorphisms at I777, indicating a low likelihood of pre-existing isolates with reduced susceptibility to ODV. Structural analysis of the RSV L polymerase active site shows I777 is in close proximity to the pre-incorporated active NTP metabolite of ODV and its parent nucleoside, but not in direct contact (Figure 5). Interestingly, I777 is part of the conserved _775_GG<u>x</u>xG_779_ sequence located within the polymerase structural motif B, which has been implicated in stabilizing the RNA template, facilitating incorporation of incoming nucleotides and translocation of nascent RNA in related viral polymerases^37,38^. The I777 amino acid is also in contact with motif F (via V614), which interacts with incoming NTP and the RNA template base^38–41^. We posit that I777L interacts with V614 to regulate positioning of F629, resulting in the modest decrease in ODV susceptibility, following a similar mechanism of action previously described for the Ebola polymerase^42^ (Figure 5B).

**Figure 5.**
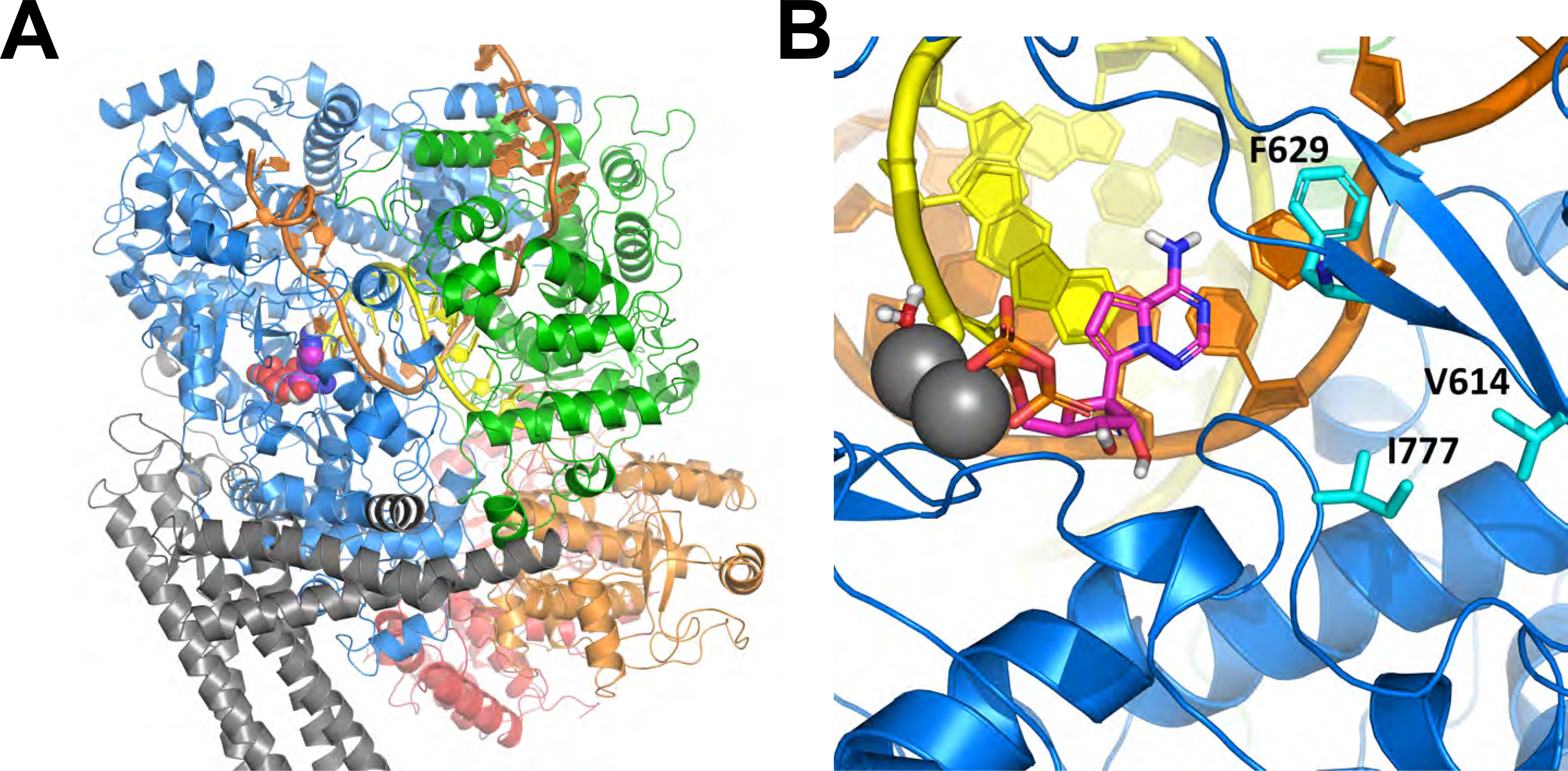
Structural analysis of I777L within the RSV polymerase active site. (A) Model of RSV L RNA-dependent RNA polymerase (RdRp) in its elongating state, with pre-incorporated GS-443902 NTP. The RdRp domain is shown in blue, the capping domain in green, the connector domain in orange, and the methyltransferase and carboxy-terminal domain (CTD) in red. The template RNA strand is in orange and the nascent RNA strand is in yellow. The phosphoprotein (P) is in grey. (B) Detail of the RSV L RdRp active site with pre-incorporated GS-443902 NTP (purple). I777 is located on motif B and is in direct contact with several hydrophobic residues on motif F, including V614.

RDV was efficacious against RSV, coronaviruses, filoviruses, and Nipah virus in nonhuman primate challenge studies^14,42–47^. While these data indicated the potential of RDV to treat RSV and supported its approval as a treatment for COVID-19, it is delivered by IV infusion due to its limited oral bioavailability. Subsequent SARS-CoV-2 nonhuman primate studies demonstrated that therapeutic efficacy could be achieved from high systemic exposures of GS-441524 resulting from either direct intravenous infusion of GS-441524^44^ or oral administration of GS-441524 prodrugs with high oral bioavailability such as ODV^15^. Additionally, a 10-day course of ODV in cynomolgus macaque models of Sudan ebolavirus and Marburg virus not only limited viral replication, but also protected animals from lethal challenge^16,17^. Here, we extended the breadth of observed in vivo efficacy with ODV to RSV in an AGM infection model. Daily oral ODV treatment significantly reduced RSV RNA in both the upper (throat swabs) and lower (BALF samples) respiratory tracts, with 69-92% reductions in the log_10_ RNA copies/mL × day AUCs observed for both groups compared to vehicle controls. At peak viral load on day 5 post-infection, treatment with 30 mg/kg or 90 mg/kg ODV resulted in >1.8 log_10_ reduction of RSV RNA in both the upper and lower respiratory tracts. These findings are in line with viral RNA reductions observed in the BALF in AGM RSV efficacy studies for lumicitabine^26^, which translated to efficacy in human RSV challenge^5^. The antiviral efficacy observed with daily ODV administration at 30 mg/kg (in a solution formulation) is notable, as this regimen in AGMs yields plasma GS-441524 exposures of approximately half that observed following a 350 mg twice daily ODV dose regimen associated with antiviral efficacy in the COVID-19 OAKTREE Phase 3 study^15,34^ (NCT05715528).

In conclusion, ODV and its parent nucleoside displayed potent antiviral activity across a wide range of RSV clinical isolates and exhibited a high barrier to resistance in vitro. Once daily oral ODV treatment was efficacious in a non-human primate model of RSV infection, significantly reducing viral RNA in both the upper and lower respiratory tracts. These data, combined with the ODV safety and tolerability data from prior COVID-19 clinical studies^32^, supported evaluation of ODV in Phase 2 clinical studies in children and high-risk adults with RSV infection (NCT06784973, NCT06585150).

## METHODS

### Compounds

Obeldesivir (ODV; GS-5245), GS-441524, ALS-8112 (lumicitabine parent nucleoside)^26^, and presatovir (GS-5806)^21,48^ were synthesized by the Department of Medicinal Chemistry, Gilead Sciences, Inc. (Foster City, CA). Sisunatovir^49^ and PC786^50,51^ were synthesized by GVK Biosciences (Hyderabad, India). Ziresovir^52^ was synthesized by Aarna Bio (Uttarakhand, India). EIDD-1931 (molnupiravir parent nucleoside)^53^ was synthesized at Enamine (Kyiv, Ukraine). Zelicapavir (EDP-938)^54^was synthesized at WuXi Apptec (Shanghai, China).

All compounds were stored frozen at −20°C in 100% dimethyl sulfoxide (DMSO) and used between 1-3 freeze-thaws.

### Cell lines

The HEp-2 cervical carcinoma cell line was purchased from American Type Culture Collection (ATCC Cat # CCL-23) and maintained at 37°C and 5% CO_2_ in Dulbecco’s Modified Eagle Medium (DMEM) + GlutaMAX (Gibco) supplemented with 10% heat-inactivated fetal bovine serum (FBS) and a 1% (v/v) of a solution containing penicillin/streptomycin (10,000 units/mL and 10,000 μg/mL; Gibco). Cells were passaged twice per week to maintain sub-confluent densities and used for experiments between passages 3-27. BHK-21 baby hamster kidney cells stably expressing the T7 polymerase (BSR-T7/5 cells) were acquired from Karl-Klaus Conzelman (Pettenkofer Institut, Munich, Germany) and maintained in Glasgow’s modified Eagle’s medium (GMEM) (Gibco) supplemented with 3% heat-inactivated FBS and 1× MEM non-essential amino acids (Gibco). BSR-T7/5 cells were grown in media supplemented with 1 mg/mL Geneticin (Gibco) every other passage to select for T7 polymerase expressing cells.

### Viruses and plasmids

Respiratory syncytial virus laboratory strain A2 was purchased from ATTC (Cat # VR-1540). The RSV A2 stock used for the inoculum for AGM efficacy studies was produced and titrated by Microbiologics. Respiratory syncytial virus B1 (Cat # NR-56243) was obtained through BEI Resources, NIAID, NIH. Clinical isolates from the United States were obtained from Pedro A. Piedra (Baylor College of Medicine, Houston, TX)^19,55,56^. Clinical isolates from the United Kingdom were obtained from Geoffrey Toms (Newcastle University, Newcastle upon Tyne, England)^19^. Belgian clinical isolates were obtained from Peter Delputte (Laboratory of Microbiology, Parasitology and Hygiene, Antwerp, Belgium)^57^. Nomenclature and GenBank accession numbers for all RSV clinical isolates utilized in these studies are listed in Supplementary Tables 1 and 2.

Plasmids used to generate RSV recombinant viruses by reverse genetics were obtained from Richard K. Plemper (Georgia State University, Atlanta, GA). These plasmids include the bacterial artificial chromosome (pBAC) containing the antigenome of RSV A2-line19F, the L shuttle vector containing the L polymerase gene, and four helper pcDNA3.1 plasmids encoding codon-optimized L polymerase, N, P, and M2-1 proteins. Both the pBAC RSV A2-line19F plasmid and the L shuttle vector contain the L polymerase gene between the MluI and SalI restriction enzyme cut sites^58–60^.

### Virus propagations

RSV laboratory, clinical isolates and recombinant RSV stocks were prepared using HEp-2 cells according to the following protocol. The day prior to infection, 2-4×10^6^ HEp-2 cells were seeded in T75 flasks and maintained overnight at 37°C and 5% CO_2_. Twenty-four hours after cell seeding, cells were washed once with 10 mL of Dulbecco’s phosphate buffered saline (DPBS) and inoculated with 7 mL total serum free DMEM containing RSV (infected at an MOI of 0.01-0.1 if titer known). Virus was incubated on cells for 2-3 hours at 37°C and 5% CO_2_, viral inoculum was then removed and replenished with 7 mL fresh DMEM supplemented with 2% heat-inactivated FBS and placed back in incubator. Viral supernatant and scraped cells were collected when substantial syncytia/cytopathic effect (CPE) were observed, and 30-50% of cells remained adherent (3-7 days post-infection). Supernatants containing cells were centrifuged for 10 minutes at 2-8 °C at 500 × *g*. Centrifuged cells were then separated from supernatant, sonicated, and recombined with supernatant. The final viral preparation was stabilized with final 20% sucrose (v/v), aliquoted, flash-frozen and stored at −80°C.

Pooled RSV A2 and clinical isolate stocks following compound selection were prepared in HEp-2 cells according to the following protocol. One day prior to infection, 7×10^5^ HEp-2 cells were seeded in T25 flasks in a final volume of 3 mL DMEM supplemented with 10% FBS. The next day, culture media was replaced with 2.7 mL DMEM supplemented with 2% heat-inactivated and 300 μL of pooled virus from a resistance selection was used to infect HEp-2 cells. To maintain substitutions already present in the RSV A2 ODV resistance selection pools, compound was added to media at the EC_90_ concentration. RSV A2 GS-441524 resistance selection pools were propagated in DMSO as growth at the EC_90_ concentration did not yield sufficient titers. Propagation of clinical RSV A and B isolate virus pools was attempted under multiple media conditions (without compound, at the compound EC_50_, or at the final selection concentration). Infected cells were assessed for levels of CPE daily and collected as described above.

### Resistance selection

Resistance selections with the RSV A2 laboratory strain, HRSV/A/Texas.USA/79254/2004, and HRSV/B/Texas.USA/79362/2005 were performed with ODV and GS-441524, as well as the fusion (F) inhibitor presatovir and a DMSO control. For all selections, HEp-2 cells were seeded in 12-well plates (one plate per compound selection) at 7.7×10^4^ cells/well in 1.5 mL DMEM supplemented with GlutaMAX and 10% heat-inactivated FBS and maintained at 37°C and 5% CO_2_ overnight. The initial infection was performed with an MOI of 0.05 in a final volume of 1.5 mL DMEM + GlutaMAX supplemented with 2% heat-inactivated FBS and compound. Cells were incubated in the presence of 1×, 2×, 5×, 10×, and 20× EC_50_ for each compound. Cultures were checked daily for the formation of CPE/syncytia and passaged when ≥50% syncytia was observed or 6-7 days after passage. Depending on levels of CPE/syncytia observed, the next passage was inoculated with 30-300 μL of cell-suspension.

### Generation of recombinant RSV A2 viruses

The reverse genetics system to generate recombinant RSV A2 has been previously described^58–61^. Briefly, the L polymerase shuttle vector was used as a template to synthesize two amplicons bearing the C319R, I777L, or L1453W substitution and homology to the MluI and SalI restriction-enzyme-digested pBAC (Supplementary Table 8). These two fragments were then cloned into the digested pBAC containing RSV antigenome by 3-fragment Gibson Assembly or In-Fusion cloning. Primer design and cloning steps were carried out in accordance with the NEBuilder HiFi DNA Assembly Master Mix (New England Biolabs). MluI and SalI restriction enzyme digests were performed following manufacturer protocols (New England Biolabs). Newly constructed recombinant pBACs bearing C319R, I777L, or L1453W were Sanger sequence-verified and transfected into BSR-T7/5 cells in combination with the four helper plasmids (L, N, P, M2-1) in a 4:1:2:2:2 ratio, respectively with Lipofectamine 3000 (Invitrogen). Transfected BSR-T7/5 cells were then continuously passaged every 2-3 days until CPE was visible. Recombinant virus was then rescued, propagated, and titered as described in Hotard et al 2012 and the virus propagation section above^58^.

### RSV indirect immunofluorescence RSV titration

HEp-2 cells were seeded 8,000 cells/well in 100 μL DMEM supplemented with 10% heat-inactivated FBS in black clear-bottom 96-well plates (Corning) and incubated 18-24 h at 37°C and 5% CO_2_. The next day, virus was serially diluted in DMEM supplemented with 2% heat-inactivated FBS and added to cells (100 μL/well). Cells were then incubated for 72 h at 37°C and 5% CO_2_. The cells were then fixed for 10 minutes at 4°C with methanol (250 μL/well) and dried for 5-10 minutes. Fixed cells were washed once with DPBS and blocked with 60 μL/well BlockAid (Invitrogen) for 30-50 minutes at room temperature. Fixed cells were then incubated with 50 μL/well of primary anti-RSV F Mab858 (Millipore) diluted 1:2000 in SuperBlock (ThermoFisher Scientific) for 1 h at room temperature. Cells were washed three times with 250 μL/well of DPBS 0.05% Tween. Fixed cells were then stained with secondary goat anti-mouse AlexaFluor647 (Invitrogen) diluted 1:2000 in SuperBlock and Hoechst (diluted 1:1000; Invitrogen) for 1 h at 37°C. The wash step was repeated and a final 100 μL/well DPBS was added. Plates were sealed with PerkinElmer black plate seal and read on a Cellomics CellInsight machine (CellHealthProfiling.V4 assay). Titers were calculated by the 50% tissue culture infectious dose (TCID_50_) Reed & Muench method using values from %High_TargetAvgIntCh2 parameter (also referred to as % infectivity), where wells were scored as positive if % infectivity was ≥ [% infectivity of negative control + (3× standard deviation of negative controls)].

### RSV antiviral assays

HEp-2 cells were seeded into black clear-bottom 96-well plates at 8,000 cells/well in 100 μL DMEM supplemented with 10% heat-inactivated FBS and incubated overnight at 37°C and 5% CO_2_. After this incubation period, media was aspirated and replenished with 100 μL/well of DMEM supplemented with 2% heat-inactivated FBS. A compound dispenser was then used to add compounds to each plate in 1:3 serial-dilutions over 8 wells starting at 0.010-50 μM concentration (set to final volume 200 μL, DMSO limit 1%). Cells were then infected with RSV at an MOI of 0.05 (in 100 μL/well; final volume in well was 200 μL). At 72 hours post-infection (hpi), cells were fixed for 10 minutes at 4°C with methanol (250 μL/well) and then dried for 5-10 minutes. Fixed cells were then washed once with DPBS and blocked with 60 μL/well BlockAid for 30-50 minutes at room temperature. Fixed cells were then incubated with 50 μL/well of primary anti-RSV F Mab858 diluted 1:2000 in SuperBlock for 1 h at room temperature. Cells were washed three times with 250 μL/well of DPBS 0.05% Tween. Fixed cells were then stained with secondary goat anti-mouse AlexaFluor647 diluted 1:2000 in SuperBlock and Hoechst (diluted 1:1000) for 1 h at 37°C. The wash step was repeated and a final 100 μL/well DPBS was added. Plates were sealed with PerkinElmer black plate seal and read on a CellInsight CX7 (CellHealthProfiling.V4 assay)^18^.

MeanTargetAvgIntensityCh2 parameter values were normalized to the uninfected and infected DMSO controls (0% and 100% infection, respectively) and plotted using log (inhibitor/agonist) vs. response (FindECanything/Variable slope – Four parameters) non-linear regression fit (GraphPad Prism 10) to determine EC_50_ values. Unless otherwise noted, all EC_50_ values were calculated as the average of at least 2-4 independent experiments and are presented as the mean ± standard deviation.

### HEp-2 cytotoxicity assays

Compounds (200 nL/well) were pre-spotted onto Grenier black 384-well plates prior to seeding HEp-2 cells (1,000 cells/well in 40 μL/well in DMEM supplemented with Glutamine, 10% heat-inactivated FBS, and Pen/Strep). Plates were incubated at 37°C and 5% CO_2_ for 72 h. After the incubation period, 20 μL/well of medium was removed and replaced with 20 μL/well of CellTiter-Glo reagent (Promega) using a Biotek washer and dispenser. Plates were read using an EnVision (PerkinElmer) with a luminescence program for 384-well plates with a 0.1 second integration time. Values were normalized to the puromycin- and DMSO-treated controls (100% and 0% cytotoxicity, respectively). Data was analyzed using a non-linear regression analysis and CC_50_ values were determined as the concentration reducing the luciferase signal by 50%. Compiled CC_50_ data were generated from n=16 biological replicates (with four technical replicates each).

### RSV RNA isolation and deep sequencing

Viral RNA was isolated from RSV clinical isolates using the MagMax Viral RNA Isolation kit (ThermoFisher) by processing 200 μL of virus sample and eluting in 100 μL nuclease-free water. Isolated RNA was used for RNA-Seq for RSV whole genome sequencing and amplicon sequencing of the RSV L gene. For RNA-Seq, isolated RNA was used for sequencing library preparation using SMARTer® Stranded Total RNA-Seq Kit v3 - Pico Input Mammalian (Takara Bio) and deep sequencing was performed using Illumina NovaSeq6000 or NextSeq2000. For RSV L amplicon-seq, cDNA synthesis was performed using the LunaScript RT SuperMix Kit (New England Biolabs) and used to PCR amplify (Platinum SuperFi PCR Master Mix Invitrogen) the RSV L gene in 9 overlapping fragments (Supplementary Table 8). The following PCR cycling parameters were used: 98°C, 30 s; followed by 30 cycles at 98°C, 20 s; 55°C, 30 s; 72°C 1 min; followed by 72°C, 5 min (Applied Biosytems Verti 96-well Thermal cycler). PCR amplicons were then pooled for each sample and the Nextera XT DNA Library Preparation Kit (Illumina) was used for preparing sequencing libraries. Deep sequencing was performed using the Illumina NextSeq2000. The RSV L amplicon-seq data was used in combination with total RNA-Seq data for two RSV clinical isolates that had low viral load or mixtures (RSV_2018_013 & RSV_2018_001). RSV L amplicon-seq data was used when available to determine changes in the L polymerase from reference.

### Data analysis of deep sequencing data

Internally developed software was used to process and align paired-end FASTQ files via a multistep method (Supplementary Table 9). Reads are aligned to an RSV subgroup reference sequence (subgroup A or B) using SMALT aligner^62^. Any read ends that overlapped with genomic coordinates of amplification primers were clipped. Tabulated nucleotides at each position were used to generate a consensus sequence. Nucleotide mixtures were reported in the consensus sequence when more than one base was present at or above 15% of the viral population. Indels were reported in the consensus sequence when present at or above 50% of the viral population. Amino acid substitution calling was tabulated based on observed codons in reads. To report in-frame insertions and deletions (indels), amino acid realignment of reads with indels was performed. For amino acid variant calling, reads containing frameshift indels were trimmed to exclude the region with an indel from further analysis. All aligned reads were then translated in-frame and changes from reference sequence were tabulated. Amino acid substitutions and indels were reported as a change from reference when present at or above 15%. For RNA-Seq, the additional steps R1-R4 were added to filter human and ribosomal reads without losing reads that are RSV (Supplementary Table 9).

### RSV lineage determination

The lineage classification system of the RSV Genotyping Consensus Consortium was implemented^63^. The Nextclade online tool (https://clades.nextstrain.org/)^64^ was used to determine lineages for RSV clinical isolates using whole genome sequences. To assign lineages, Nextclade incorporates sequences on a phylogenetic reference tree based on curated RSV-A and RSV-B sequences. Phylogenetic trees were visualized with Nextstrain Auspice (https://docs.nextstrain.org/projects/auspice/en/stable/).

### Determination of sequence conservation and selected substitution polymorphisms

RSV sequences and metadata were downloaded from nextstrain.org/rsv on 12 December 2023. High quality sequences (qc.overallStatus = good) with more than 90% genome coverage were included and viral sequences from non-human hosts were removed. This resulted in a dataset of 2891 RSV A sequences and 2181 RSV B sequences with broad geographic distribution (Supplementary Table 10). The RSV L polymerase sequence was extracted from downloaded sequences and then translated to amino acid sequences. Amino acid sequences were then aligned to L polymerase amino acid sequences of references NC_038235 (RSV A2) or NC_001781 (RSV B1). Amino acid conservation was calculated as the frequency of the most common amino acid at each position in coordinates of the corresponding subgroup reference sequence. Conservation analysis was performed for RSV A and RSV B separately. To identify compound-selected substitutions, amino acid variations in the RSV L polymerase during passages with compound were compared with the viral stock RSV L polymerase sequence (passage [P]0). Substitutions in the L polymerase at ≥15% prevalence in ≥2 successive passages and not identified in the DMSO passages were reported. Conservation at these amino acids was assessed as described above for public RSV sequences. A residue with amino acid conservation <99% was considered a polymorphism.

### Generation of RSV L polymerase structure model

Several structures of the RSV A2 L polymerase in complex with the phosphoprotein (P) have been determined with cryo-electron microscopy^38,65–67^. The full-length L polymerase is 2165 amino acids, but none of these structures resolves residues beyond ∼1460 amino acids. While the RdRp and capping domains are clearly resolved, the connector domain (CD), the methytransferase (MTase), and the C-terminal domain (CTD) are not. We predicted these domains using AlphaFold 2.1^68^, and the domains were combined with the cryo-EM structure 6PZK^38^ to produce a more complete model of the RSV L polymerase. The general arrangement of the domains followed the conformation observed in the rabies L polymerase structure 6UEB^69^. From this starting point, models of the RNA initiation and elongation states were created, based on structures for influenza B (6QCT)^70^ and other viral L polymerases such as VSV (5A22)^71^. All models were refined using software available in the Schrӧdinger Suite (Schrödinger Release 2024-2: Schrödinger, LLC, New York, 2024). Although the RSV B L polymerase structure remains unsolved, RSV A and B L polymerases share ∼90% amino acid sequence identity and are predicted to be structurally homologous^72^.

### Animal care

The in vivo efficacy study was conducted at Lovelace Biomedical Research Institute (LBRI, Albuquerque, NM). All study protocols were reviewed by the Institutional Animal Care and Use Committee (IACUC) at LBRI. All studies were conducted under an IACUC approved protocol in compliance with the Animal Welfare Act, Public Health Service policy, and other federal statutes and regulations relating to animals and experiments involving research animals. Research facilities at LBRI are accredited by the Association for Assessment and Accreditation of Laboratory Animal Care, International and strictly adhere to principles stated in the Guide for Care and Use of Laboratory Animals, National Research Council, 2011 (National Academies Press, Washington, DC). The efficacy study, which involved animals experimentally infected with RSV, was conducted in an animal biosafety level 2 (ABSL-2) laboratory.

Wild-caught African green monkeys (AGMs; *Chlorocebus aethiops*), of St. Kitts origin, were sourced through Worldwide Primates Inc. (Florida, USA). AGMs were housed in adjacent individual cages, within a climate-controlled room with a fixed light/dark cycle (12-h light/12-h dark). AGMs were monitored at least twice daily for the duration of the study by trained personnel. Commercial monkey chow, treats and fruit were provided twice daily. Water was available to the AGMs *ad libitum*. Prior to initiation of the study, AGMs were acclimated to pole-collar capture and chair restraint to facilitate oral dosing by oral gavage on days that did not coincide with other sample collection (i.e., days 2, 4 and 6 post-challenge).

### Reagents and formulations for in vivo studies

ODV was diluted into the vehicle control solution (2.5% dimethyl sulfoxide (DMSO), 2.5% propylene glycol, 10% Labrasol, 10% Kolliphor HS-15, and 75% water) at concentrations of 15 mg/mL (30 mg/kg dose group) or 45 mg/mL (90 mg/kg dose group).

### AGM RSV antiviral efficacy study

Fifteen AGMs (7 male/8 female equally distributed across dosing groups) were utilized to compare the efficacy of orally delivered ODV at 30 or 90 mg/kg in the RSV model of infection. On study day 0, animals were anesthetized with ketamine (10 mg/kg) followed by inhaled isoflurane (up to 5% in oxygen) for RSV infection. Animals received a total inoculum of ∼2.6 x 10^6^ TCID_50_ RSV A2 by combined intranasal (0.5 mL in each nostril) and intratracheal (1 mL) instillation. Approximately 20-28 hours post RSV infection, animals were dosed with ODV or vehicle control administered via oral gavage at a volume of 2 mL/kg. Treatment was continued once daily for a total of 6 days. On treatment days that coincided with collection of throat swabs and BALF samples (i.e., days 1, 3 and 5 post-challenge), animals were dosed while anesthetized, immediately following sample collection. On the remaining treatment days (i.e., days 2, 4 and 6 post-challenge), dosing was performed in alert, chair-restrained animals.

Throat swabs were collected at baseline, day 1 post-infection, and every other day through day 15 post-infection using a cotton-tipped applicator presoaked in sterile saline. Swabs were placed in a tube containing 0.5 mL sterile saline, frozen immediately on dry ice, and stored at −70°C until further processing for analysis by real-time quantitative PCR (RT-qPCR).

Bronchioalveolar lavage fluid (BALF) was collected from left and right caudal lung lobes at baseline, day 1 post-infection, and every other day through day 15 post-infection. For BALF collection, a pediatric bronchoscope (Olympus XP-40) was advanced into the caudal lung lobe, 10 mL of sterile saline was infused, and the maximum volume was aspirated. BALF was centrifuged at 1000 ×*g* for 10 minutes at 4°C. The resulting cell pellet and 1 mL aliquots of supernatant were flash frozen and stored at −70°C until further processing for analysis by RT-qPCR.

### RNA isolation and RT-qPCR

RT-qPCR was performed on RNA extracted from BALF supernatant and throat swabs collected from each animal at each timepoint. The RNA from 100 uL BALF sample was extracted using the MagMAX Pathogen RNA isolation kit (Thermofisher) on the KingFisher Flex (ThermoFisher) and performed according to manufacturer instructions, including the addition of the exogenous RNA extraction control (VetMAX Xeno Internal Positive Control RNA (ThermoFisher)). The extraction buffer used for sample extraction was spiked with Xeno RNA and the combined extraction buffer was mixed with the BALF immediately after thawing. The Xeno control RNA served as an extraction efficiency and matrix inhibition control to account for loss or inhibition of target RNA during downstream RT-qPCR analysis. The isolated RNA was eluted in 75 uL (swabs) or 80 uL (BALF) RNase free water. RSV RNA loads were then measured by RT-qPCR for the RSV N gene. Copies per mL equivalents were calculated from a standard curve generated from RSV DNA plasmid stocks of known copy concentration. A set of RSV DNA plasmid serial dilutions to generate standard curve were added for each RT-qPCR run. RSV N gene concentrations for each sample were presented as copies/mL equivalents based on the standard curve run with each reaction. All samples were run in duplicate, for BALF samples the values provided are an average of the left and right lung samples. A fixed volume of RNA (5 μL) was added to a reaction mix of 1× TaqMan® Fast Virus 1-Step Master Mix (ThermoFisher), 1× RSV N gene Primer Probe Mix (containing 900 nM of primers and 250 nM of probe) and water was added to make up a final volume of 20 μL. The RSV N gene primers and probe sequences are listed in Supplementary Table 8.

In a separate reaction for the internal positive control the 1× Xeno™ VIC™ Primer Probe Mix replaced the RSV primers. Amplification and detection were performed using a QuantStudio 7 Flex Thermal Cycler under the following cycling conditions: 50 °C for 5 min, 95 °C for 20 seconds and 40 cycles of 95 °C for 15 seconds, 60 °C for 60 seconds.

### Statistical analysis

All statistics were performed using GraphPad Prism 10.1.2. For AGM efficacy studies, longitudinal analyses of the ODV 30 or 90 mg/kg treatments were compared to the vehicle control group by repeated measures two-way ANOVA with Bonferroni post-hoc correction for multiple comparisons. Samples analyzed by RT-qPCR that were detectable but below the lower limit of quantification (LLOQ) for the assay were assigned a value of ½ of the LLOQ, while undetectable samples were assigned a value of ¼ the LLOQ before conversion to log scale. The LLOQs for these analyses are 3.3 and 2.3 log_10_ RNA copies/mL for the throat swab and BALF, respectively. AUC analyses were conducted with the baseline set at the LLOQ for each sample type for each animal and taking the mean of each group. For statistical analysis of AUC, a one-way ANOVA with Bonferroni post-hoc correction for multiple comparisons was utilized. Corrected *p* values of <0.05 for were considered statistically significant for all analyses conducted.

## ACKNOWLEDGEMENTS

We thank Lovelace Biomedical support teams for contributions to the execution of the RSV AGM efficacy model, Karl-Klaus Conzelman (Pettenkofer Institut, Munich, Germany) for the BSR-T7/5 cells, Richard K. Plemper (Georgia State University, Atlanta, GA) for recombinant RSV plasmids, Geoffrey Toms (formerly of Newcastle University, Newcastle upon Tyne, England) for UK RSV isolates.

## AUTHOR CONTRIBUTIONS

JP and JLRZ contributed equally to this work and are listed by alphabetical order. Author contributions are as follows. Conceptualization: JP, JLRZ, DB, RS, RLM, TC, SPF, JPB.; Methodology: JLRZ, JP, SM, TA, AV, MSV, CR, DH, JKP, KHS.; Data curation: JP, JLRZ, TA, RM, AC.; Formal Analysis: JP, JLRZ SM, TA.; Investigation: JLRZ, SM, TA, JC, VC, VN, NCR, AV, MSV, SE, CR, DH, SC, IL, CM, NP, KHS, AC; Project administration: RM, CH, DSS, CM, RM, JS, HM, RS, RLM, TC, SPF, JPB.; Resources: JLRZ, TA, RM, JM, RK, VA, PAP, KS, PLD; Supervision: JP, JLRZ, SM, TA, MSV, KHS, RM, JM, CH, DSS, CM, RM, JS, HM, RS, RLM, TC, SPF, JPB.; Validation: JLRZ, SM, TA, JC, VC, VN, NCR, MSV, SE, CR, DH, SC, IL, CM, NP, AC.; Visualization: JLRZ, JP, SM, TA, JKP.; Writing-original draft: JP, JLZR, SM, TA, JC, JPB; Writing-review and editing: all authors.

