## Supplemental Information for "Oral dosing of the nucleoside analog obeldesivir is efficacious against RSV infection in African green monkeys"

**Supplementary Information**

**Supplementary Table 1.** ODV and GS-441524 antiviral activity against RSV A clinical isolates
**Supplementary Table 2.** ODV and GS-441524 antiviral activity against RSV B clinical isolates
**Supplementary Table 3.** Substitutions in RSV L polymerase and F protein during selection with ODV, GS-441524, or presatovir
**Supplementary Table 4.** Phenotypic characterization of presatovir-passaged RSV virus pools
**Supplementary Table 5.** Phenotypic analysis of ODV- and GS-441524-selected RSV A2 virus pools
**Supplementary Table 6.** Cross-resistance phenotype of recombinant RSV A2 bearing L polymerase substitutions C319R, I777L, or L1453W
**Supplementary Table 7.** RSV RNA loads in throat swabs and BALF
**Supplementary Table 8.** Primers used in this study
**Supplementary Table 9.** Overview of deep sequencing analysis pipeline for Illumina
**Supplementary Table 10.** Geographic origin of sequences included in the dataset
**Supplementary Figure 1.** ODV and GS-441524 potencies against RSV A and B clinical isolates
**Supplementary Figure 2.** RSV A and B clinical isolate phylogeny trees
**Supplementary Figure 3.** RSV L polymerase structural analysis of ODV and GS-441524 selected substitutions

**Supplementary Table 1. ODV and GS-441524 antiviral activity against RSV A clinical isolates.**

| RSV A strain or isolate[a] | GenBank ID | Lineage[b] | ODV | | GS-441524 | |
|---|---|---|---|---|---|---|
| | | | $EC_{50}$ (µM)[c] | Fold change from RSV A2[d] | $EC_{50}$ (µM)[c] | Fold change from RSV A2[d] |
| RSV A2 (LS) | NC_038235 | A.1 | 0.46 ± 0.16 | 1.00 | 1.00 ± 0.35 | 1.00 |
| HRSV/A/Texas.USA/A1-37425/1987 | PQ416565 | A | 0.35 ± 0.22 | 0.76 | 0.42 ± 0.44 | 0.42 |
| HRSV/A/Texas.USA/A4-66125/1992 | PQ416566 | A.2 | 0.34 ± 0.07 | 0.74 | 0.78 ± 0.11 | 0.78 |
| HRSV/A/Texas.USA/61245/1994 | PQ416562 | A.2 | 0.25 ± 0.12 | 0.54 | 0.56 ± 0.13 | 0.56 |
| HRSV/A/Texas.USA/79254/2004 | PQ416563 | A.2.1 | 0.56 ± 0.16 | 1.22 | 1.04 ± 0.28 | 1.04 |
| HRSV/A/Texas.USA/79256/2004 | PQ416564 | A.3 | 0.27 ± 0.07 | 0.59 | 0.32 ± 0.03 | 0.32 |
| HRSV/A/Texas.USA/79303/2005 | OK649678 | A.3 | 0.30 ± 0.08 | 0.65 | 0.74 ± 0.29 | 0.74 |
| HRSV/A/Texas.USA/79312/2005 | PQ416561 | A.3 | 0.34 ± 0.13 | 0.74 | 0.57 ± 0.10 | 0.57 |
| HRSV/A/GBR/NU5108/2008 | PQ591394 | A.3.1 | 0.33 ± 0.06 | 0.72 | 0.77 ± 0.28 | 0.77 |
| HRSV/A/GBR/NU4908/2008 | PQ591395 | A.3.1 | 0.55 ± 0.21 | 1.20 | 1.49 ± 0.31 | 1.49 |
| HRSV/A/GBR/NU4408/2008 | PQ591393 | A.2.1 | 0.34 ± 0.18 | 0.74 | 0.75 ± 0.32 | 0.75 |
| HRSV/A/GBR/NU0409/2009 | PQ591396 | A.2.1.1 | 0.52 ± 0.22 | 1.13 | 1.21 ± 0.27 | 1.21 |
| HRSV/A/BEL/RVRA013/2018 | PQ376590 | A.D.3 | 0.39 ± 0.08 | 0.85 | 0.90 ± 0.32 | 0.90 |
| HRSV/A/BEL/RVRA080/2019 | PQ376589 | A.D.2.2 | 0.60 ± 0.15 | 1.30 | 0.85 ± 0.28 | 0.85 |
| HRSV/A/USA/TX-HOU-R06687/2021 | PQ308736 | A.D.1 | 0.39 ± 0.20 | 0.85 | 1.03 ± 0.21 | 1.03 |
| HRSV/A/BEL/RVRA126/2020 | PQ376588 | A.D.5.2 | 0.35 ± 0.10 | 0.76 | 0.66 ± 0.24 | 0.66 |
| HRSV/A/BEL/RVRA011/2023 | PQ376586 | A.D.1 | 0.53 ± 0.07 | 1.15 | 0.88 ± 0.12 | 0.88 |
| HRSV/A/BEL/RVRA012/2023 | PQ376587 | A.D.1 | 0.31 ± 0.22 | 0.67 | 0.70 ± 0.49 | 0.70 |
| Mean of all RSV A isolates ± SD | | | 0.40 ± 0.11 | 0.86 ± 0.24 | 0.80 ± 0.28 | 0.80 ± 0.28 |
| Range of all RSV A isolates (min – max) | | | 0.25 – 0.60 | 0.55 – 1.32 | 0.32 – 1.49 | 0.32 – 1.49 |
| $CC_{50}$ (µM) in HEp-2 cells | | | >50[e] | | >100[f] | |

ODV = obeldesivir; LS = laboratory strain; $EC_{50}$ = 50% effective concentration; $CC_{50}$ = 50% cytotoxic concentration

[a] RSV clinical isolates named following CDC universal nomenclature: HRSV/subgroup/geographic identifier/unique sequence identifier/year of sampling as previously described[1].

[b] Lineages are characterized as previously described [2]. [c] The data represent the mean ± standard deviation (SD) of at least two independent experiments. [d] Fold change from RSV A2 values were calculated by dividing each virus isolate mean $EC_{50}$ by the mean $EC_{50}$ of RSV A2. [e] $CC_{50}$ values represent data from n=16 experiments. [f] $CC_{50}$ value previously reported[3].

### Supplementary Table 2. ODV and GS-441524 antiviral activity against RSV B clinical isolates

| RSV B strain or isolate[a] | GenBank ID | Lineage[b] | ODV | | GS-441524 | |
|---|---|---|---|---|---|---|
| | | | $EC_{50}$ (µM)[c] | Fold change from RSV A2[d] | $EC_{50}$ (µM)[c] | Fold change from RSV A2[d] |
| RSV A2 (LS) | NC_038235 | A.1 | 0.46 ± 0.16 | 1.00 | 1.00 ± 0.35 | 1.00 |
| RSV B1 (LS) | NC_001781 | B.2 | 0.43 ± 0.07 | 0.93 | 0.91 ± 0.12 | 0.91 |
| HRSV/B/Texas.USA/60823/1993 | PQ416569 | B | 0.36± 0.15 | 0.78 | 0.76 ± 0.10 | 0.76 |
| HRSV/B/Texas.USA/60687/1993 | PQ416570 | B | 0.30 ± 0.16 | 0.65 | 0.63 ± 0.46 | 0.63 |
| HRSV/B/Texas.USA/61501/1993 | PQ416568 | B | 0.62 ± 0.08 | 1.35 | 1.24 ± 0.07 | 1.24 |
| HRSV/B/GBR/NU1705/1996 | PQ591399 | B.3 | 0.45 ± 0.23 | 0.98 | 1.45 ± 0.25 | 1.45 |
| HRSV/B/GBR/NU613/1997 | PQ591398 | B | 0.27 ± 0.18 | 0.59 | 0.59 ± 0.26 | 0.59 |
| HRSV/B/Texas.USA/79233/2004 | PQ416567 | B.D | 0.20 ± 0.09 | 0.43 | 0.55 ± 0.40 | 0.55 |
| HRSV/B/Texas.USA/79362/2005 | OK649754 | B.D | 0.47 ± 0.17 | 1.02 | 0.96 ± 0.12 | 0.96 |
| HRSV/B/GBR/NU0609/2009 | PQ591397 | B.D | 0.52 ± 0.20 | 1.13 | 0.95 ± 0.36 | 0.95 |
| HRSV/B/USA/TX-HOU-813046/2015 | PQ308737 | B.D.4.1 | 0.65 ± 0.26 | 1.41 | 1.59 ± 0.53 | 1.59 |
| HRSV/B/BEL/RVRA013/2017 | PQ376592 | B.D.4.1 | 0.29 ± 0.03 | 0.63 | 0.75 ± 0.33 | 0.75 |
| HRSV/B/USA/TX-HOU-IP0035B/2018 | PQ308738 | B.D.4.1.1 | 0.35 ± 0.08 | 0.76 | 0.83 ± 0.35 | 0.83 |
| HRSV/B/BEL/RVRA001/2018 | PQ376591 | B.D.4.1.1 | 0.66 ± 0.21 | 1.43 | 1.18 ± 0.10 | 1.18 |
| HRSV/B/BEL/RVRA107/2019 | PQ376594 | B.D.4.1.1 | 0.49 ± 0.10 | 1.07 | 0.97 ± 0.08 | 0.97 |
| HRSV/B/USA/TX-HOU-IP0187A/2021 | PQ308739 | B.D.E.1 | 0.62 ± 0.09 | 1.35 | 1.18 ± 0.28 | 1.18 |
| HRSV/B/BEL/RVRA002/2022 | PQ376593 | B.D.4.1.1 | 0.35 ± 0.03 | 0.76 | 0.86 ± 0.00 | 0.86 |
| Mean of all RSV B isolates ± SD | | | 0.44 ± 0.15 | 0.95 ± 0.32 | 0.96 ± 0.30 | 0.96 ± 0.30 |
| Range of all RSV B isolates (min – max) | | | 0.20 – 0.66 | 0.43 – 1.43 | 0.55 – 1.59 | 0.55 – 1.59 |

ODV = obeldesivir; LS = laboratory strain; $EC_{50}$ = 50% effective concentration

[a] RSV clinical isolates named following CDC universal nomenclature: HRSV/subgroup/geographic identifier/unique sequence identifier/year of sampling as previously described[1].

[b] Lineages are characterized as previously described [2]. [c] The data represent the mean ± standard deviation (SD) of at least two independent experiments. [d] Fold change from RSV A2 values were calculated by dividing each virus isolate mean $EC_{50}$ by the mean $EC_{50}$ of RSV A2.

**Supplementary Table 3. Substitutions in RSV L polymerase or F protein during selection with ODV, GS-441524, or presatovir**

| RSV strain | L Polymerase Substitutions Detected[a] (Passages Observed) | | F Substitutions Detected[b] (Passages Observed) |
|---|---|---|---|
| | ODV selection | GS-441524 selection | Presatovir selection |
| RSV A2 | **I777I/L** (P7)<br>**I777L** (P8-P13) | **L1453L/W** (P5-P13) | **F140F/I** (P3-P4)<br>**P320P/H** (P3-P4) |
| HRSV/A/Texas.USA/79254/2004 | **K1188K/E** (P2-P3) | - | **D486N** (P1-P2) |
| HRSV/B/Texas.USA/79362/2005 | - | **C319C/R** (P4-P10)<br>**C319R** (P11-P13)<br>**C1881C/Y** (P4-P5)<br>**S169S/P** (P5-P9)<br>**T660T/A** (P5-P8) | **M396M/K** (P3-P4)<br>**E487E/V** (P3-P4) |

[a]Sequences were determined by L amplicon next-generation sequencing. [b]Sequences were determined by next-generation sequencing of total RNA. All substitutions noted are those that were detected in the L polymerase or F protein at ≥15% of sequencing reads for ≥2 successive passages and were not observed in DMSO control passaged virus. P = passage

### Supplementary Table 4. Phenotypic characterization of presatovir-passaged RSV virus pools

| Compound | Inhibitor class | RSV A2 | | | HRSV/A/Texas.USA/79254/2004 | | | HRSV/B/Texas.USA/79362/2005 | | |
|---|---|---|---|---|---|---|---|---|---|---|
| | | Input (P0) | Presatovir-selected | | Input (P0) | Presatovir-selected | | Input (P0) | Presatovir-selected | |
| | | EC50 (μM)[a] | EC50 (μM)[a] | Fold change from P0[b] | EC50 (μM)[a] | EC50 (μM)[a] | Fold change from P0[b] | EC50 (μM)[a] | EC50 (μM)[a] | Fold change from P0[b] |
| Obeldesivir | Nuc | 0.53 ± 0.09 | 0.45 ± 0.16 | 0.89 ± 0.34 | 0.91 ± 0.12 | 0.42 ± 0.13 | 0.47 ± 0.15 | 1.01 ± 0.19 | 0.69 ± 0.19 | 0.67 ± 0.06 |
| GS-441524 | Nuc | 0.93 ± 0.22 | 0.57 ± 0.31 | 0.57 ± 0.20 | 1.25 ± 0.41 | 0.68 ± 0.27 | 0.53 ± 0.04 | 0.95 ± 0.22 | 0.71 ± 0.23 | 0.73 ± 0.07 |
| Remdesivir | Nuc | 0.019 ± 0.004 | 0.013 ± 0.001 | 0.73 ± 0.21 | 0.023 ± 0.004 | 0.013 ± 0.000 | 0.58 ± 0.001 | 0.026 ± 0.009 | 0.016 ± 0.004 | 0.65 ± 0.06 |
| Zelicapavir (EDP-938)[4] | N-inhibitor | 0.062 ± 0.00005 | 0.062 ± 0.004 | 1.01 ± 0.07 | 0.059 ± 0.004 | 0.049 ± 0.004 | 0.84 ± 0.13 | 0.088 ± 0.013 | 0.068 ± 0.008 | 0.78 ± 0.02 |
| Presatovir[5] | F-inhibitor | 0.000051 ± 0.000007 | UND | UND | 0.00038 ± 0.00012 | 0.0022 ± 0.0003 | 190 ± 190 | 0.00044 ± 0.00022 | UND | UND |
| Sisunatovir[6] | F-inhibitor | 0.000046 ± 0.000017 | UND | UND | 0.00075 ± 0.00036 | 0.0061 ± 0.0060 | 5.63 ± 5.28 | 0.00096 ± 0.00035 | UND | UND |

Nuc = nucleoside inhibitor, F= fusion/entry, N = nucleoprotein, NNI = non-nucleoside viral polymerase inhibitor, UND = undetermined - $EC_{50}$ could not be determined as inhibition never reached 50%, thus fold-change also cannot be determined.

[a] The data represent the mean ± standard deviation (SD) of two independent experiments

[b] Fold-change calculated by ($EC_{50}$ of pool)/($EC_{50}$ of input RSV A2 virus) from individual replicates then taking the average ± SD from fold-change of replicates

### Supplementary Table 5. Phenotypic analysis of ODV- and GS-441524-selected RSV A2 virus pools

| Compound | Inhibitor class | RSV A2 Input (P0) | DMSO-passaged RSV A2 | | ODV-selected pool (I777L RSV A2) | | GS-441524-selected pool (L1453L/W RSV A2) | |
|---|---|---|---|---|---|---|---|---|
| | | $EC_{50}$ (µM)[a] | $EC_{50}$ (µM)[a] | Fold change from P0[b] | $EC_{50}$ (µM)[a] | Fold change from P0[b] | $EC_{50}$ (µM)[a] | Fold change from P0[b] |
| Obeldesivir | Nuc | 0.71 ± 0.39 | 0.44 ± 0.26 | 0.65 ± 0.20 | 1.67 ± 0.69 | 2.82 ± 0.84 | 0.86 ± 0.31 | 1.53 ± 0.58 |
| GS-441524 | Nuc | 1.41 ± 0.50 | 0.84 ± 0.33 | 0.59 ± 0.14 | 3.04 ± 0.41 | 2.37 ± 0.59 | 1.74 ± 0.53 | 1.27 ± 0.20 |
| Remdesivir | Nuc | 0.026 ± 0.012 | 0.017 ± 0.003 | 0.71 ± 0.20 | 0.084 ± 0.028 | 3.36 ± 0.77 | 0.026 ± 0.005 | 1.12 ± 0.38 |
| Lumicitabine (ALS-8112)[7] | Nuc | 3.54 ± 0.68 | 2.57 ± 0.09 | 0.75 ± 0.13 | 3.31 ± 0.87 | 0.93 ± 0.18 | 4.49 ± 1.96 | 1.21 ± 0.33 |
| Molnupiravir (EIDD-1931)[8] | Nuc | 8.37 ± 4.56 | 4.82 ± 2.55 | 0.64 ± 0.18 | 1.69 ± 1.03 | 0.19 ± 0.03 | 7.52 ± 3.60 | 0.99 ± 0.19 |
| Zelicapavir (EDP-938)[4] | N-inhibitor | 0.057 ± 0.009 | 0.040 ± 0.006 | 0.72 ± 0.17 | 0.049 ± 0.005 | 0.87 ± 0.06 | 0.042 ± 0.003 | 0.75 ± 0.13 |
| Presatovir[5] | F-inhibitor | 0.00025 ± 0.00011 | 0.00011 ± 0.00003 | 0.56 ± 0.23 | 0.00014 ± 0.00001 | 0.71 ± 0.35 | 0.00014 ± 0.00005 | 0.61 ± 0.10 |
| Sisunatovir[6] | F-inhibitor | 0.00014 ± 0.00007 | 0.000065 ± 0.000024 | 0.56 ± 0.18 | 0.000076 ± 0.000012 | 0.82 ± 0.54 | 0.000087 ± 0.000033 | 0.79 ± 0.31 |
| PC786[9] | NNI | 0.0011 ± 0.0004 | 0.00088 ± 0.00031 | 0.86 ± 0.06 | 0.0012 ± 0.0003 | 1.24 ± 0.24 | 0.0012 ± 0.0004 | 1.21 ± 0.26 |

ODV = obeldesivir; $EC_{50}$ = 50% effective concentration; Nuc = nucleoside inhibitor, F = fusion/entry, N = nucleoprotein, NNI = non-nucleoside viral polymerase inhibitor
[a] The data represent the mean ± standard deviation (SD) of three independent experiments performed in triplicate
[b] Fold-change calculated by ($EC_{50}$ of pool)/($EC_{50}$ of input RSV A2 virus) from individual replicates ± SD

**Supplementary Table 6. Cross-resistance phenotype of recombinant RSV A2 bearing L polymerase substitutions C319R, I777L, or L1453W**

| Compound | Inhibitor class | Wildtype rRSV A2 reference $EC_{50}$ (µM)[a] | rRSV A2 C319R $EC_{50}$ (µM)[a] | Fold change from rRSV A2[b] | Wildtype rRSV A2 reference $EC_{50}$ (µM)[a] | rRSV A2 I777L $EC_{50}$ (µM)[a] | Fold change from rRSV A2[b] | Wildtype rRSV A2 reference $EC_{50}$ (µM)[a] | rRSV A2 L1453W $EC_{50}$ (µM)[a] | Fold change from rRSV A2[b] |
|---|---|---|---|---|---|---|---|---|---|---|
| ODV | Nuc | 0.43 ± 0.11 | 0.73 ± 0.01 | 1.83 ± 0.45 | 0.39 ± 0.04 | 1.51 ± 0.47 | 3.83 ± 0.86 | 0.26 ± 0.04 | 0.27 ± 0.11 | 1.06 ± 0.50 |
| GS-441524 | Nuc | 0.60 ± 0.11 | 1.01 ± 0.13 | 1.72 ± 0.52 | 0.70 ± 0.20 | 2.30 ± 0.49 | 3.31 ± 0.25 | 0.79 ± 0.11 | 0.74 ± 0.25 | 0.95 ± 0.39 |
| Remdesivir | Nuc | 0.020 ± 0.001 | 0.027 ± 0.000 | 1.39 ± 0.07 | 0.012 ± 0.001 | 0.033 ± 0.010 | 2.73 ± 0.66 | 0.016 ± 0.003 | 0.015 ± 0.002 | 0.99 ± 0.17 |
| Lumicitabine[7] (ALS-8112) | Nuc | 0.93 ± 0.13 | 2.03 ± 0.03 | 2.21 ± 0.35 | 0.91 ± 0.05 | 0.89 ± 0.32 | 0.97 ± 0.30 | 0.56 ± 0.14 | 0.72 ± 0.22 | 1.28 ± 0.18 |
| Molnupiravir[8] (EIDD-1931) | Nuc | 2.94 ± 0.30 | 4.53 ± 0.22 | 1.55 ± 0.23 | 3.74 ± 1.00 | 0.98 ± 0.63 | 0.25 ± 0.10 | 2.16 ± 0.26 | 3.27 ± 0.50 | 1.54 ± 0.42 |
| Zelicapavir[4] (EDP-938) | N-inhibitor | 0.071 ± 0.003 | 0.090 ± 0.007 | 1.27 ± 0.15 | 0.059 ± 0.002 | 0.045 ± 0.005 | 0.77 ± 0.12 | 0.048 ± 0.006 | 0.052 ± 0.013 | 1.07 ± 0.20 |
| Presatovir[5] | F-inhibitor | 0.00012 ± 0.00006 | 0.00019 ± 0.00011 | 2.07 ± 1.91 | 0.000037 ± 0.000019 | 0.000066 ± 0.000025 | 2.29 ± 1.87 | 0.000054 ± 0.000011 | 0.000063 ± 0.000011 | 1.18 ± 0.10 |
| Sisunatovir[6] | F-inhibitor | 0.000061 ± 0.000030 | 0.00014 ± 0.00016 | 2.61 ± 1.02 | 0.000029 ± 0.000001 | 0.000024 ± 0.000009 | 0.83 ± 0.30 | 0.000039 ± 0.000012 | 0.000030 ± 0.000004 | 0.83 ± 0.25 |
| PC786[9] | NNI | 0.00045 ± 0.00002 | 0.00085 ± 0.00004 | 1.89 ± 0.16 | 0.00029 ± 0.00003 | 0.00030 ± 0.00004 | 1.06 ± 0.24 | 0.00024 ± 0.00006 | 0.00051 ± 0.00007 | 2.22 ± 0.54 |

ODV = obeldesivir; $EC_{50}$ = 50% effective concentration; Nuc = nucleoside inhibitor; F = fusion/entry; N = nucleoprotein; NNI = non-nucleoside L polymerase inhibitor; rRSV = recombinant RSV

[a] The data represent the mean ± standard deviation of at least two independent experiments

[b] $EC_{50}$ fold change values were calculated by dividing each virus $EC_{50}$ by the $EC_{50}$ of the wildtype RSV reference strain that was tested concurrently with rRSV C319R, I777L or L1453W for each individual replicate

### Supplementary Table 7. RSV RNA loads in throat swabs and BALF.

| | | Days post infection: | Baseline | 1 | 3 | 5 | 7 | 9 | 11 | 13 | 15 | AUC |
|---|---|---|---|---|---|---|---|---|---|---|---|---|
| Sample | Group | RSV A2 RNA load ($\log_{10}$ copies/mL) | | | | | | | | | | RNA $\log_{10}$ copies/mL · day |
| Throat Swabs | Vehicle | Mean | 3.27 | 3.57 | 5.51 | 6.26 | 6.10 | 4.60 | 3.57 | 3.34 | 3.33 | 13.63 |
| | | SD | 0.00 | 0.00 | 0.29 | 0.56 | 0.47 | 0.98 | 0.00 | 0.10 | 0.14 | 2.80 |
| | PO ODV 30 mg/kg | Mean | 3.33 | 3.45 | 3.69 | 4.46 | 4.84 | 4.39 | 3.55 | 3.34 | 3.34 | 4.26 |
| | | SD | 0.14 | 0.17 | 0.45 | 0.62 | 0.79 | 0.84 | 0.06 | 0.10 | 0.10 | 0.74 |
| | | *p* | 0.999 | 0.979 | <0.0001 | <0.0001 | <0.0001 | 0.853 | >0.999 | >0.999 | >0.999 | <0.0001 |
| | PO ODV 90 mg/kg | Mean | 3.27 | 3.46 | 3.31 | 3.51 | 3.60 | 3.45 | 4.03 | 3.88 | 3.46 | 1.05 |
| | | SD | 0.00 | 0.12 | 0.08 | 0.38 | 0.57 | 0.17 | 0.57 | 0.55 | 0.12 | 0.61 |
| | | *p* | >0.999 | 0.985 | <0.0001 | <0.0001 | <0.0001 | 0.0002 | 0.303 | 0.169 | 0.972 | <0.0001 |
| BALF | Vehicle | Mean | 2.30 | 3.01 | 5.43 | 5.81 | 5.63 | 3.85 | 2.56 | 2.30 | 2.30 | 18.29 |
| | | SD | 0.00 | 0.60 | 0.23 | 0.24 | 0.51 | 0.40 | 0.49 | 0.00 | 0.00 | 2.03 |
| | PO ODV 30 mg/kg | Mean | 2.36 | 2.61 | 3.18 | 3.77 | 3.04 | 3.45 | 2.47 | 2.30 | 2.30 | 3.96 |
| | | SD | 0.08 | 0.26 | 0.42 | 0.74 | 0.65 | 0.67 | 0.11 | 0.00 | 0.00 | 2.92 |
| | | *p* | 0.999 | 0.383 | <0.0001 | <0.0001 | <0.0001 | 0.385 | 0.990 | >0.999 | >0.999 | <0.0001 |
| | PO ODV 90 mg/kg | Mean | 2.30 | 2.74 | 2.90 | 3.18 | 2.86 | 2.56 | 2.49 | 2.39 | 2.30 | 1.83 |
| | | SD | 0.00 | 0.53 | 0.55 | 0.67 | 0.73 | 0.18 | 0.12 | 0.09 | 0.00 | 2.51 |
| | | *p* | >0.999 | 0.702 | <0.0001 | <0.0001 | <0.0001 | <0.0001 | 0.997 | 0.992 | >0.999 | <0.0001 |

ODV = obeldesivir; BALF = bronchoalveolar lavage fluid; SD = standard deviation

RSV RNA copies from throat swabs and BALF collected on odd days between day 1 and day 15 post-infection from AGMs (African green monkeys; n=5/group) dosed daily with PO vehicle, 30 mg/kg ODV, or 90 mg/kg ODV. The lower limit of quantification for these assays were 7500 copies/mL and 800 copies/mL for throat swabs and BALF, respectively. Samples below the lower limit of quantitation (LLOQ) were assigned a value of ½ LLOQ, while samples with undetectable levels of RNA were assigned a value of ¼ LLOQ. All values were then log transformed for statistical analyses. Group means, standard deviation, and *p* values compared to vehicle control at each timepoint using repeated measures two-way ANOVA with Bonferroni post-hoc correction are presented. Area under the curve (AUC), using the LLOQ as baseline, was determined for each individual animal and the group mean, standard deviation, and *p* values compared to vehicle control from one-way ANOVA with Bonferroni post-hoc correction. For all statistical analyses, $p < 0.05$ is considered significant.

### Supplementary Table 8. Primers used in this study.

| Name | (5'-to-3') | Description |
|---|---|---|
| Forward_Primer_Fragment 1 | ataaaacacaattgaatgccagtcgaccttaccatctg | In-Fusion Cloning Primer for C319R |
| Reverse_Primer_C319R_Fragment 1 | gctttagtatacgatctccataaagg | In-Fusion Cloning Primer for C319R |
| Forward_Primer_C319R_Fragment 2 | cctttatggagatcgtatactaaagc | In-Fusion Cloning Primer for C319R |
| Reverse_Primer_Fragment 2 | aatgggtcgagaagcttacgcgtatatagttcctcctttc | In-Fusion Cloning Primer for C319R |
| Forward_Primer_Fragment 1 | tatagttatataaaacacaattgaatgccagtcgaccttaccatctgt | Gibson Assembly Primer for I777L |
| Reverse_Primer_I777L_Fragment 1 | cagtttttgacaccacccttccaggccacccatgtgatatctatataatc | Gibson Assembly Primer for I777L |
| Forward_Primer_I777L_Fragment 2 | gattatatagatatcacatgggtggcctggaagggtggtgtcaaaaactg | Gibson Assembly Primer for I777L |
| Reverse_Primer_Fragment 2 | gaatgggtcgagaagcttacgcgtatatagttcctcctttcagc | Gibson Assembly Primer for I777L |
| Forward_Primer_Fragment 1 | tatagttatataaaacacaattgaatgccagtcgaccttaccatctgt | Gibson Assembly Primer for L1453W |
| Reverse_Primer_L1453W_Fragment 1 | ccacatattgagtccaacttattttgtctgg | Gibson Assembly Primer for L1453W |
| Forward_Primer_L1453W_Fragment 2 | ccagacaaaataagttggactcaatatgtgg | Gibson Assembly Primer for L1453W |
| Reverse_Primer_Fragment 2 | gaatgggtcgagaagcttacgcgtatatagttcctcctttcagc | Gibson Assembly Primer for L1453W |
| Forward_Primer_RSV A2 N | gctcttagcaaagtcaagttgaatga | RSV N gene RT-qPCR Primer |
| Reverse_Primer_RSV A2 N | tgctccgttggatggtgtatt | RSV N gene RT-qPCR Primer |
| RSV A2 Probe | 6FAM-acactcaacaaagatcaacttctgtcatccagc-MGBNFQ | RSV N gene RT-qPCR Probe |
| RSV-L1for | taagagtgtacaatactgt | RSV L amplicon 1 |
| RSV-L1rev | tattaaccatgatggaggatg | RSV L amplicon 1 |
| RSV-L2for | gctatagaaathagtgatgt | RSV L amplicon 2 |
| RSV-L2rev | acagcatccatkgcttgtct | RSV L amplicon 2 |
| RSV-L3for | agcttgcaggtgayaataa | RSV L amplicon 3 |
| RSV-L3rev | tctagatcaccatatcttgt | RSV L amplicon 3 |
| RSV-L4for | agaatgtttgcwatgcaacc | RSV L amplicon 4 |
| RSV-L4rev | ctaaatattaaactgcataata | RSV L amplicon 4 |
| RSV-L5for | catgctcaagcagattatt | RSV L amplicon 5 |
| RSV-L5rev | gtatatttratgtccattg | RSV L amplicon 5 |
| RSV-L6for | tggtcwttatccaatatagt | RSV L amplicon 6 |

| | | |
|---|---|---|
| RSV-L6rev | ccagctaaattagtactta | RSV L amplicon 6 |
| RSV-L7for | tgagatacatttgatgaa | RSV L amplicon 7 |
| RSV-L7rev | cacctatgaatgctataca | RSV L amplicon 7 |
| RSV-L8for | tgcatgcttccttggcatc | RSV L amplicon 8 |
| RSV-L8rev | acttcattacgtccagctatag | RSV L amplicon 8 |
| RSV-L9for | gtttacttagtccttacaatag | RSV L amplicon 9 |
| RSVA-L9rev | tctcgtagtttagttaata | RSV L amplicon 9 |
| RSVB-L9rev | tctcgttgtgttgtaaatg | RSV L amplicon 9 |

**Supplementary Table 9. Overview of deep sequencing analysis pipeline for Illumina.**

| Step | Description |
|---|---|
| R1 | Build combined reference index for GCA_000001405.15 GRCH38, and RSV-A (RSV_A_NC_038235) and RSV-B (NC_001781) references using BWA[10]<br>Align FASTQ files to combined index using BWA aligner to split out any reads aligning to host. Generate FASTQ files containing viral and unmapped reads from BWA alignment |
| R2 | Filter out additional host reads using SortMeRNA[11]. Settings: --sam – num_alignments 1 –fastx and with the following ribosomal reference databases:<br>silva-bac-16s-id90.fasta, silva-bac-23s-id98.fasta, silva-arc-16s-id95.fasta, silva-arc-23s-id98.fasta, silva-euk-18s-id95.fasta, silva-euk-28s-id98.fasta, rfam-5s-database-id98.fasta, rfam-5.8s-database-id98.fasta |
| R3 | Estimate the percentage of RSV reads by aligning 100,000 reads against the RSV subtype references and counting the number of aligned reads |
| R4 | Use Seqtk (https://github.com/lh3/seqtk) to reduce the number of paired reads to keep an average read depth of 20,000 if more RSV reads than that are available. Command:<br>seqtk sample-s 123 \|fastq_file\| num_reads<br>with num_reads calculated based on the desired read coverage, read length and genome length. |
| 1 | Trim and filter low quality reads using Trimmomatic[12]. Settings: SLIDINGWINDOW:4:15 MINLEN: 50. In addition, clip any adapters using ILLUMINACLIP: bbmap adapters: fa:2:30:10 |
| 2 | Contigs are generated from paired-end FASTQ files by VICUNA assembler[13] per gene. Blast contigs against a curated set of RSV subtype A and B references to determine subtype<br>For amplicon (L): Identify subtype, and chose subtype reference<br>For RNA-seq (full genome): Identify the closest subtype match from a curated set of sequences. Use this subtype reference to guide de-novo assembly from contigs |
| 3 | Merge trimmed, paired-end reads that overlap using Ngmerge[14]. Create FASTQ file containing merged reads and any single end reads that do not overlap |
| 4 | Align trimmed, merged reads to corresponding RSV subtype reference (NC_038235 for subtype A and NC_001781 for subtype B, for L) or de-novo assembly (for full genome) using SMALT aligner[15] |
| 5 | If applicable, clip amplification primers from aligned reads based on genomic coordinates |
| 6 | Filter out reads with length smaller 50 bp or low quality (average phred score <30) |
| 7 | Tabulate nucleotide variants per position in target gene coordinates of corresponding RSV subtype reference |

| | |
|---|---|
| 8 | Generate consensus sequences with 15% cutoff and include ambiguity codes for any positions with more than one nucleotide exceeding cutoff. Indels are included in consensus if they exceed 50% frequency<br><br>a. The read depth at each position must exceed 50 reads to report variant or indel, otherwise an ambiguous base (N) is reported in consensus sequence<br>b. Exclude nucleotide variants with low average phred score (<20) |
| 9 | Tabulate mutations at amino acid level in gene regions<br><br>a. If read contains indel(s), evaluate read for number of insertions and deletions. If sum of insertions minus sum of deletions is a multiple of 3, perform amino acid alignment to realign indels. If not multiple of 3, extract longest fragment between two indels or 5' end and indel or indel and 3' end<br>b. Rollup amino acid variants and indels across amplification pools per sample.<br>c. Exclude any amino acid variant with low average phred score (<20)<br>d. Report amino acid variants and indels at or above 15% cutoff<br>e. Exclude any amino acid variants or indels at gene positions with read depth less than 50 reads<br>f. Map all variant calls to positions of genotype-specific reference sequence. |

**Supplementary Table 10. Geographic origin of sequences included in the dataset**

| Region/ Country# | RSV-A (N=2891) (% of all RSV-A) | RSV-B (N=2181) (% of all RSV-B) | RSV-A and RSV-B (N=5072) |
|---|---|---|---|
| Africa | 518 (17.9%) | 536 (24.6%) | 1054 (20.8%) |
| Asia | 265 (9.2%) | 163 (7.4%) | 428 (8.5%) |
| Europe | 335 (11.6%) | 338 (15.5%) | 673 (13.3%) |
| North America | 1174 (40.6%) | 606 (27.8%) | 1780 (35.1%) |
| Oceania | 258 (8.9%) | 250 (11.5%) | 508 (10.0%) |
| South America | 315 (10.9%) | 219 (10.0%) | 534 (10.6%) |
| Unknown | 26 (0.9%) | 69 (3.1%) | 95 (1.9%) |
| Total all Regions | 2891 | 2181 | 5072 |

#Africa (Kenya, South Africa, Uganda, Zambia), Asia (China, Hong Kong, India, Japan, Jordan, Lebanon, Philippines, South Korea, Thailand, Vietnam), Europe (Austria, Belgium, France, Germany, Italy, Netherlands, Russia, Spain, Switzerland, United Kingdom), North America (Canada, Mexico, USA, Nicaragua), Oceania (Australia, New Zealand), South America (Argentina, Brazil, Peru)

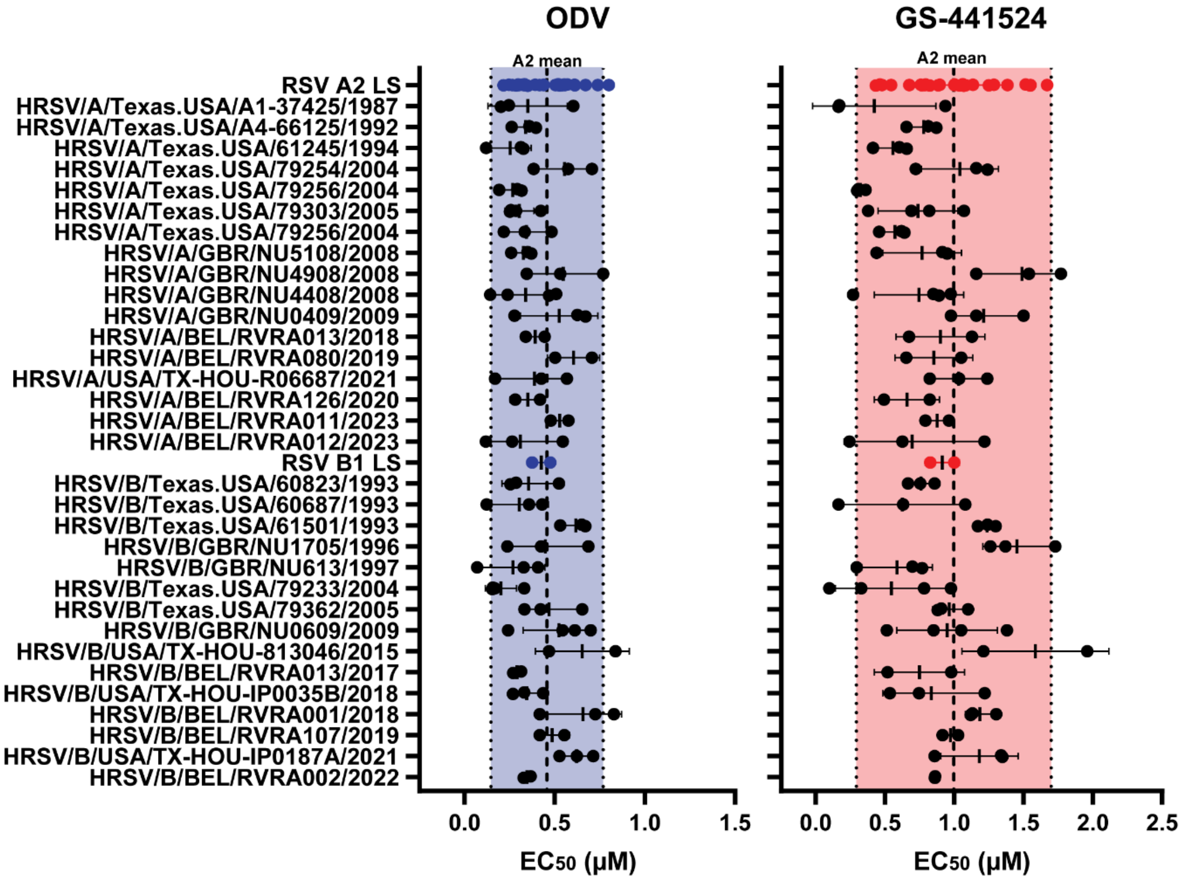


**Supplementary Figure 1. ODV and GS-441524 potencies against RSV A and B clinical isolates.** Half-maximal effective concentration ($EC_{50}$) values of ODV (shaded in blue) and GS-441524 (shaded in red) against the RSV A2 laboratory strain (RSV A2 LS), the RSV B1 laboratory strain (RSV B1 LS), 17 RSV A clinical isolates, and 15 RSV B clinical isolates. Each data point represents a biological replicate, with the mean (vertical tick) and standard deviation (error bars). The dashed line represents the RSV A2 mean of all biological replicates and the dotted lines represent plus or minus 2× the standard deviation of RSV A2 mean $EC_{50}$.

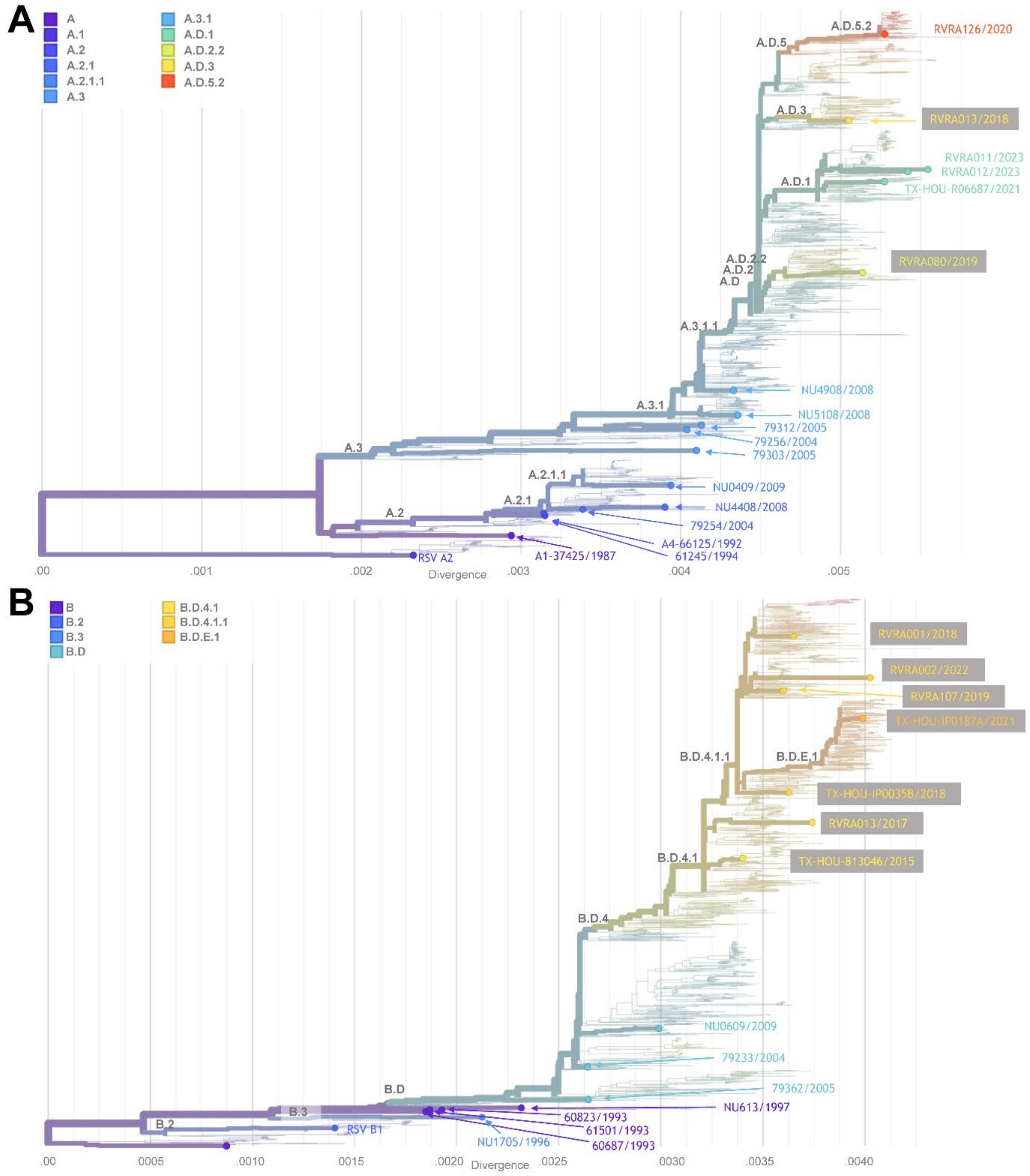


**Supplementary Figure 2. RSV A and B clinical isolate phylogeny trees.** Phylogeny trees generated from whole genome sequences for RSV A (A) and RSV B (B) isolates. The phylogenic trees were visualized with Nextstrain Auspice and lineages assigned using the Nextclade online tool. Isolates are indicated by the unique sequence identifier and year of sampling (from Supplementary Tables 1 and 2). X-axis shows nucleotide divergence. Lineages are characterized as previously described[2]**.**

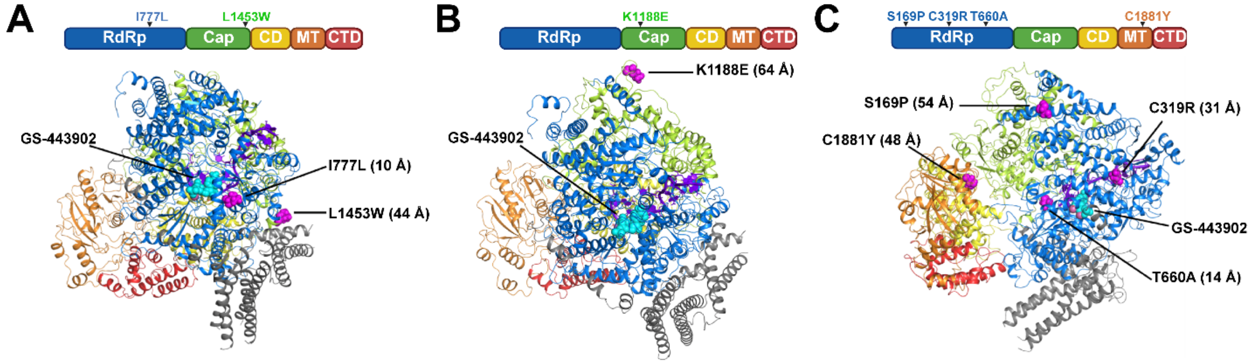


**Supplementary Figure 3**. **RSV L polymerase structural analysis of ODV- and GS-441524-selected substitutions.** Model of RSV L:P complex in its RNA initiation state with pre-incorporated GS-443902 (cyan), the active NTP metabolite of ODV and GS-441524. The L polymerase RdRp domain is in blue, the capping domain (Cap) in green, the connector domain (CD) in yellow, the methyltransferase (MT) domain in orange, and the C-terminal domain (CTD) in red. The primer and template RNA are in purple, and the tetrameric phosphoproteins are in grey. (A) RSV A2 L polymerase selected substitutions I777L and L1453W are shown in magenta. As measured from the residue Cα to the C1′ of GS-443902, I777L (located in the RdRp active site), is 10 Å from the pre-incorporated inhibitor GS-443902, and L1453W (located in the capping domain), is 44 Å from the pre-incorporated inhibitor GS-443902. (B) RSV A clinical isolate L polymerase substitution K1188E is shown in magenta located within the capping domain. The Cα of K1188E is 64 Å from the C1′ of the pre-incorporated GS-443902. (C) RSV B clinical isolate L polymerase substitutions S169P (in the RdRp domain), C319R (in the RdRp domain), T660A (in the RdRp domain) and C1881Y (in the methyltransferase domain) are shown in magenta. Measured distances from residues' Cα to the C1′ of GS-443902 are 54 Å (S169P), 31 Å (C319R), 14 Å (T660A) and 48 Å (C1881Y)
